# Escape From Oncogene-Induced Senescence is Controlled by POU2F2 and Memorized by Chromatin Scars

**DOI:** 10.1101/2022.06.29.497942

**Authors:** Ricardo Iván Martínez-Zamudio, Alketa Stefa, José Américo Nabuco, Themistoklis Vasilopoulos, Mark Simpson, Gregory Doré, Pierre-François Roux, Mark A. Galan, Ravi J. Chokshi, Oliver Bischof, Utz Herbig

## Abstract

Although oncogene-induced senescence (OIS) is a potent tumor-suppressor mechanism, recent studies revealed that cells can escape from OIS with features of transformed cells. However, the mechanisms that promote OIS escape remain unclear, and evidence of post-senescent cells in human cancers is missing. Here, we unravel the regulatory mechanisms underlying OIS escape using dynamic multidimensional profiling. We demonstrate a critical role for AP1 and POU2F2 transcription factors for escape from OIS and identify ‘senescence-associated chromatin scars (SACS)’ as an epigenetic memory of OIS, detectable during colorectal cancer progression. POU2F2 levels are elevated already in precancerous lesions and as cells escape from OIS, and its expression and binding activity to cis-regulatory elements are associated with decreased patient survival. Our results support a model in which POU2F2 exploits a precoded enhancer landscape to promote senescence escape and reveal POU2F2 gene signatures and SACS as valuable biomarkers with diagnostic and prognostic potential.

## Introduction

After a brief period of hyperproliferation, cells that overexpress specific oncogenes, such as constitutively active H-RAS^G12V^, undergo oncogene-induced senescence (OIS)^1^, a multifaceted cell state that arrests cells at risk for malignant transformation. Cells with features of OIS have been detected in early neoplastic and premalignant lesions in humans, some of which remain inactive for years ^2, 3^. However, a subset of these lesions eventually progresses to more advanced cancer stages, an event that is associated with the loss of certain senescence features ^3–6^. The evolution of premalignant lesions into more aggressive cancer stages therefore raises the possibility that senescent cells within these lesions develop mechanisms that allow them to escape from OIS. Elucidating these mechanisms can open inroads into the early detection, patient stratification, and intervention of premalignant lesions that remain susceptible to cancer progression.

Cells in OIS remain arrested due to activation of the p53/p21 and/or p16/retinoblastoma (Rb) tumor suppressor pathways ^7–10^. In addition, these senescent cells secrete a complex mixture of inflammatory cytokines, proteases, and stemness factors called the senescence-associated secretory phenotype (SASP), which further stabilizes the senescence state in an autocrine manner but also affects neighboring cells in a paracrine manner ^11, 12^. Paradoxically, the paracrine effects of the SASP have been shown to facilitate transdifferentiation ^13, 14^, reprogramming ^15^, stemness ^16^, and cancer development ^17^, demonstrating that senescent cells, through their SASP, can destabilize the identity of cells within their immediate environment.

Whether a perturbed SASP output can also destabilize the proliferative arrest of cells in OIS remains unknown.

The stability of OIS is determined by the oncogene and the cellular context under which the signaling occurs ^18, 19^. Furthermore, due to the dynamic nature of senescence ^16, 20^, cells have been shown to escape from the senescence state through cell-autonomous and cell non-autonomous mechanisms ^21–23^. Indeed, recent studies have demonstrated that cells that remained in the senescent state for prolonged periods develop means to resume proliferation and develop features of cancers cells by mechanisms that involve derepression of the *hTERT* locus ^24^, downregulation of histone demethylases ^25^, reorganization of topologically associated domains (TADs) ^26^ and stemness-associated reprogramming ^21^. However, while the molecular and phenotypic characteristics acquired by cells escaping the senescence state recapitulates the evolutionary process driving oncogenic transformation^3^, the gene regulatory mechanisms underlying senescence escape and evidence of this phenomenon in human cancer are still absent.

The epigenetic changes that lead to the activation of the senescence state have only recently been investigated from a dynamic perspective. Several studies have revealed transcription factors (TFs) and enhancers as critical determinants for the timely execution of the senescence program and for priming the senescent epigenome to respond to future incoming stimuli ^27–30^.

These studies, therefore, have shifted our view of cellular senescence from that of a static cell fate to that of a temporally organized process akin to (epi)genetically programmed differentiation and cellular responses to environmental cues. Under this paradigm, the dynamic remodeling of the epigenome, driven by interactions between TFs and enhancers throughout the different phases of senescence, predetermines the potential of senescent cells to promote distinct (patho)physiological outcomes (i.e., reprogramming, immune clearance, transformation).

Despite these advances in our understanding of the temporal evolution of the senescence response, the dynamic interplay between TFs and the enhancer landscape of senescent cells that promotes escape from OIS remains fragmentary. In light of the clinical relevance of the potential instability of OIS during cancer development, a detailed understanding of the gene regulatory mechanisms underlying this process can reveal windows of opportunity for the modulation of senescent cells in premalignant lesions that are susceptible to cancer progression.

In this study, we defined previously unknown mechanisms of senescence escape by generating and integrating time-resolved gene expression and epigenomic profiles of cells escaping from oncogenic RAS-induced senescence. This approach identified TFs and TF networks regulating senescence escape, potential targets for novel cancer therapies, and senescence-escape signatures in human colorectal cancer (CRC) with diagnostic and prognostic potential.

## Results

### DYNAMIC MULTI-OMIC PROFILING TO DECIPHER OIS ESCAPE

Previous studies, including ours, demonstrated that OIS is an unstable cell fate from which cells can escape, either by activating hTERT expression or by promoting chromosomal inversions ^24, 26^. Given the highly dynamic nature of the senescence arrest, we hypothesized that senescence escape might occur through multiple routes. Indeed, normal human primary fibroblasts (strain GM21) expressing oncogenic H-RAS^G12V^ frequently escaped from OIS between days 18-25 following transduction of the oncogene, yet their escape often appeared to be independent of telomerase activation as cells did not always retain high levels of hTERT activity or expression levels (**Figures S1A,B,J).** Despite this, cells that escaped from OIS displayed anchorage-independent growth, a hallmark of oncogenic transformation ^31, 32^, and developed abortive colonies in soft agar (**Figures S1C-F**). Consistent with this, our results revealed that H-RAS^G12V^ expressing GM21 fibroblasts that escaped from OIS eventually ceased proliferation with features of telomere dysfunction-induced senescence at around day 80 after transduction of the oncogene (**Figures S1G,H**). H-RAS^G12V^ expression levels remained elevated throughout the entire time course, with a moderate decrease of RAS levels as cells escaped from OIS, which is consistent with previous observations ^24^ (**Figure S1I**). To identify the underlying mechanism for OIS escape, we applied a time-resolved profiling approach as previously described by us ^24, 27^. We generated and integrated time-resolved bulk profiles of gene expression using microarrays, gene regulatory elements (promoters and enhancers) using ChIP-seq of histone modifications H3K27ac and -K4me1, and TF binding dynamics using ATAC-seq across four stages: proliferation (P; empty vector controls), senescence (S, days 8-17), transition (T, days 18-25) and escape (E, day 26+) (**Figure 1A**). We tracked senescence by measuring classical senescence biomarkers and features, detecting these only in the senescence and transition stages. Notably, SASP factors IL1ß and IL8 remained upregulated in post-senescent cells, although at significantly lower expression levels (**Figures S1J, K**).

**Figure 1.**
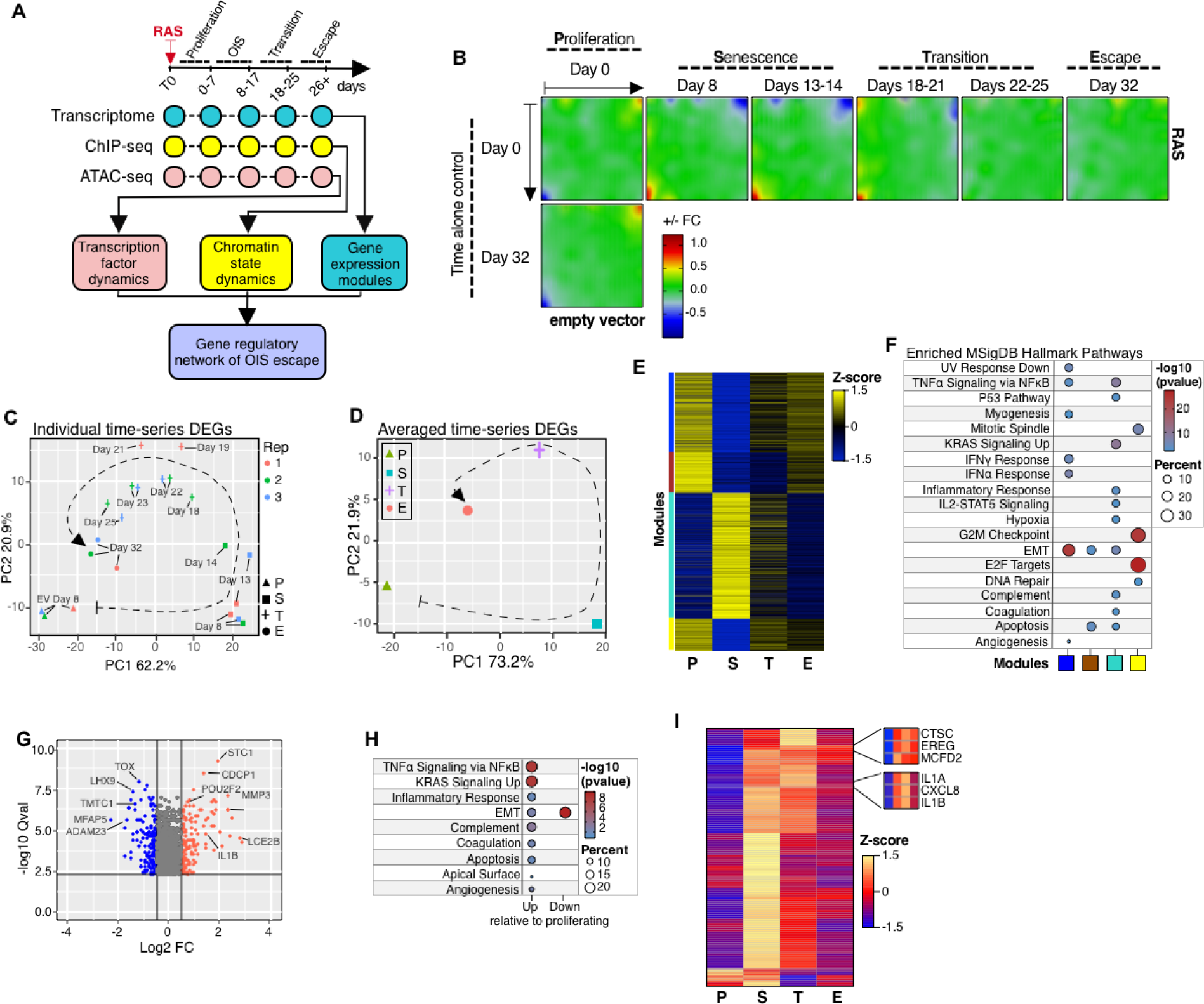
Transcriptional landscape of OIS escape. **(A)** Diagram describing the time-resolved multi-omic (transcriptome, microarray; epigenome, ChIP-seq & ATAC-seq) profiling approach used to define the gene-regulatory mechanisms of escape from OIS. **(B)**, Averaged self- organizing maps (SOMs) of transcriptomes of three biologically independent time-series experiments of GM21 fibroblasts undergoing escape from OIS (horizontal) and time-matched empty vector control (vertical) expressed as logarithmic fold-change. The transcriptional landscape of each stage of the OIS escape process is highlighted. Bold initials for each stage are used as abbreviations throughout the study. P, proliferation; S, senescence; T, transition; E, escape. **(C**,**D)** Principal component analysis projection plots showing the individual (**C**) and averaged (**D**) transcriptional trajectories of differentially expressed genes of three biologically independent OIS escape time-series experiments. **(E)** Heatmap showing the color-coded modules (blue, brown, yellow, and turquoise) of differentially expressed genes of GM21 fibroblasts in P, S, T, and E. The average of three biologically independent time-series experiments is shown. **(F)** Functional over-representation analysis map showing significant associations of the Molecular Signatures Database (MSigDB) Hallmark gene sets for each module described in (**E**). Circle fill is color-coded according to the FDR-corrected *p-value* from a hypergeometric distribution test. Circle size is proportional to the percentage of genes in each MSigDB gene set found within each gene module. *N* > 300 genes per module. **(G)** Volcano plot showing differentially expressed genes in cells after OIS escape, relative to empty vector control proliferating cells. **(H)** Functional over-representation analysis map showing significant associations of the Molecular Signatures Database (MSigDB) Hallmark gene sets with genes Up and Down in cells that escaped OIS relative to proliferating control cells. Circle fill is color- coded according to the FDR-corrected *p-value* from a hypergeometric distribution test. Circle size is proportional to the percentage of genes in each MSigDB gene set found within each gene module. Up, 118 genes; Down, 150 genes. **(I)** Heatmap showing the expression levels of SASP genes from the OIS-specific turquoise module (118 genes predicted to be secreted) of cells in P, S, T, and E. Examples are shown in insets. Data are averaged from three biologically independent time-series experiments.

### MAPPING THE TRANSCRIPTIONAL LANDSCAPE OF OIS ESCAPE

To visualize transcriptional dynamics of OIS escape, we utilized a self-organizing map (SOM) machine-learning algorithm ^33^ and principal component analysis (PCA). We used time-matched control cells transduced with an empty vector as a reference. Significantly, while the transcriptional landscape of control cells remained essentially identical until the experiments ended on day 56, the transcriptome of cells expressing H-Ras^G12V^ evolved substantially through the senescence, transition, and escape stages **(Figure 1B** and **Figure S2A)**. Transcriptional trajectories of differentially expressed genes (DEGs) and global transcriptomes confirmed these results, as revealed by PCA (**Figures 1C,D, Figure S2B**). Furthermore, the transcriptional landscape of cells that entered and escaped from OIS was distinct from that of time-matched controls, as shown by SOM portraits, D-cluster projection, and metagene expression analysis ^33^ (**Figures S2C,D;** see the behavior of clusters A, F, and K as examples).

To characterize the transcriptional evolution of OIS escape in greater detail, we performed gene clustering and pathway enrichment analyses. We identified 2,118 differentially regulated genes (DEGs) (950 up; 1,168 down), which were partitioned into four distinct modules (blue, brown, turquoise, and yellow), thereby revealing state-specific transcription patterns (**Figure 1E, Figure S2E**). Consistent with a senescence arrest, OIS was characterized by upregulation of genes related to the SASP, RAS, and P53 pathways in the turquoise module. In contrast, downregulated genes, represented by the blue, brown, and yellow modules, were enriched for cell cycle, DNA repair, and EMT pathways (**Figure 1F**). The OIS transition state was defined by gradual downregulation of SASP genes and upregulation of cell cycle genes, leading up to OIS escape. At that point, cells resumed proliferation and featured a transcriptional profile that remained markedly distinct from that of normal proliferating fibroblasts (**Figures 1E**,**F**).

Significantly, cells that escaped from OIS retained higher expression levels of a subset of senescence-associated genes involved in inflammatory signaling and cell plasticity, compared to control cells (**Figures 1G,H**), which is consistent with our RT-qPCR data (**Figure S1F**. In addition, a subset of genes encoding SASP factors, which were highly expressed during the OIS state, remained upregulated in cells that escaped OIS relative to control cells, hinting at a carry-over of a senescence signature in cells that escaped from OIS (**Figure 1I**). Importantly, up-regulated SASP factors such as CTSC, EREG, MCFD2, IL1α, IL1ß and IL8, are up-regulated in several tumor types and act as unfavorable prognostic markers ^34^. The transcriptional dynamics were faithfully reproduced in three biological replicates, suggesting that cell state succession for OIS escape is largely precoded (**Figure S2F**). Overall, these results highlight the dynamic and highly organized nature of cells’ transcriptional transitions as they enter and escape from OIS.

### SENESCENCE ENHANCER REMODELING DICTATES TRANSCRIPTIONAL TRANSITIONS AND OIS ESCAPE

Growth past senescence barriers may be a pivotal event in the earliest steps of carcinogenesis. How the epigenetic landscape dictates transcriptional transitions and creates windows of opportunity for OIS escape is an open question with important clinical implications. To address this question, we characterized genomic regulatory element (in)activation (promoters and enhancers). We interrogated TF binding dynamics using ChIP-seq profiling of histone modifications (i.e., H3K4me1 and H3K27ac) and conducted transposon-accessible chromatin profiling using ATAC-seq on control cells at day 0 (P) and on H-RAS^G12V^ expressing cells isolated on days 10 (S), 18, 22 (T), and 32 (E). ChromstaR analysis ^33^ identified eight chromatin states (**Figure S3A**). While around 82% of the genome lacked enhancer-associated histone modifications and accessible chromatin, approximately 8% of chromatin was associated with active histone marks H3K4me1 and H3K27ac, as well as with accessible DNA regions (**Figure S3B**). We quantified transitions by merging redundant chromatin states (i.e., by adding regions defined by H3K4me1/H3K27ac and regions defined by H3K4me1/H3K27ac + ATAC) involving activation and inactivation of enhancers and promoters. We identified prominent enhancer activation from the unmarked and poised states as cells entered OIS (day 10), followed by dynamic enhancer remodeling during the transition stage (days 18 and 22; **Figures 2A,B**). In contrast, promoters were only modestly remodeled (**Figures 2A**,**B**). Visualization of the signal distribution that originated from the unmarked state at day 0 revealed gradual gains of TF binding and enhancer activation as cells entered OIS, peaking at days 14-18 as the cells approached the transition stage, after which enhancers were gradually inactivated and reverted to a poised state (**Figure S3C**). Similar kinetics of enhancer activation was observed at poised enhancers (defined at day 0; **Figure S3D**). In contrast, active enhancers underwent cycles of activation and inactivation, returning to levels comparable to control cells after cells escaped from OIS (**Fig S3E**). Consistent with our published results^27^, footprinting analysis identified the AP1 TF superfamily as the key bookmarking agent of prospective senescence enhancers, defining senescence-associated transcriptional transitions (**Figures S3F,G**). In addition, TFs, including IRF1, MZF1, and FOXC2, were enriched at active enhancers, suggesting a role of these TFs in cell proliferation (**Figure S3H**). Our analysis also revealed two poorly characterized zinc finger proteins, ZNF384 and ZNF263, as factors pervasively associated with putative enhancer regions. Integrating chromatin state transitions to transcriptional output using corresponding analysis (CA) revealed a robust correlation between the chronology of chromatin state transitions with individual gene modules (**Figure 2C**). Upon examining the expression of genes proximal to eight most frequent chromatin state transitions, we observed greatest transcriptional variability during OIS (days 8 and 14, **Figure 2D**), after which the gene expression output stabilized to levels that were distinct from control cells. In contrast, the chromatin states acquired during the senescence phase were preserved generally in post-senescent cells (**Figure 2D**, top of panels). Collectively, our data suggest that dynamic enhancer remodeling predicates OIS transcriptional transitions, creates a window of opportunity for OIS escape, and lays down an epigenetic memory of senescence in cells that escaped from OIS.

**Figure 2.**
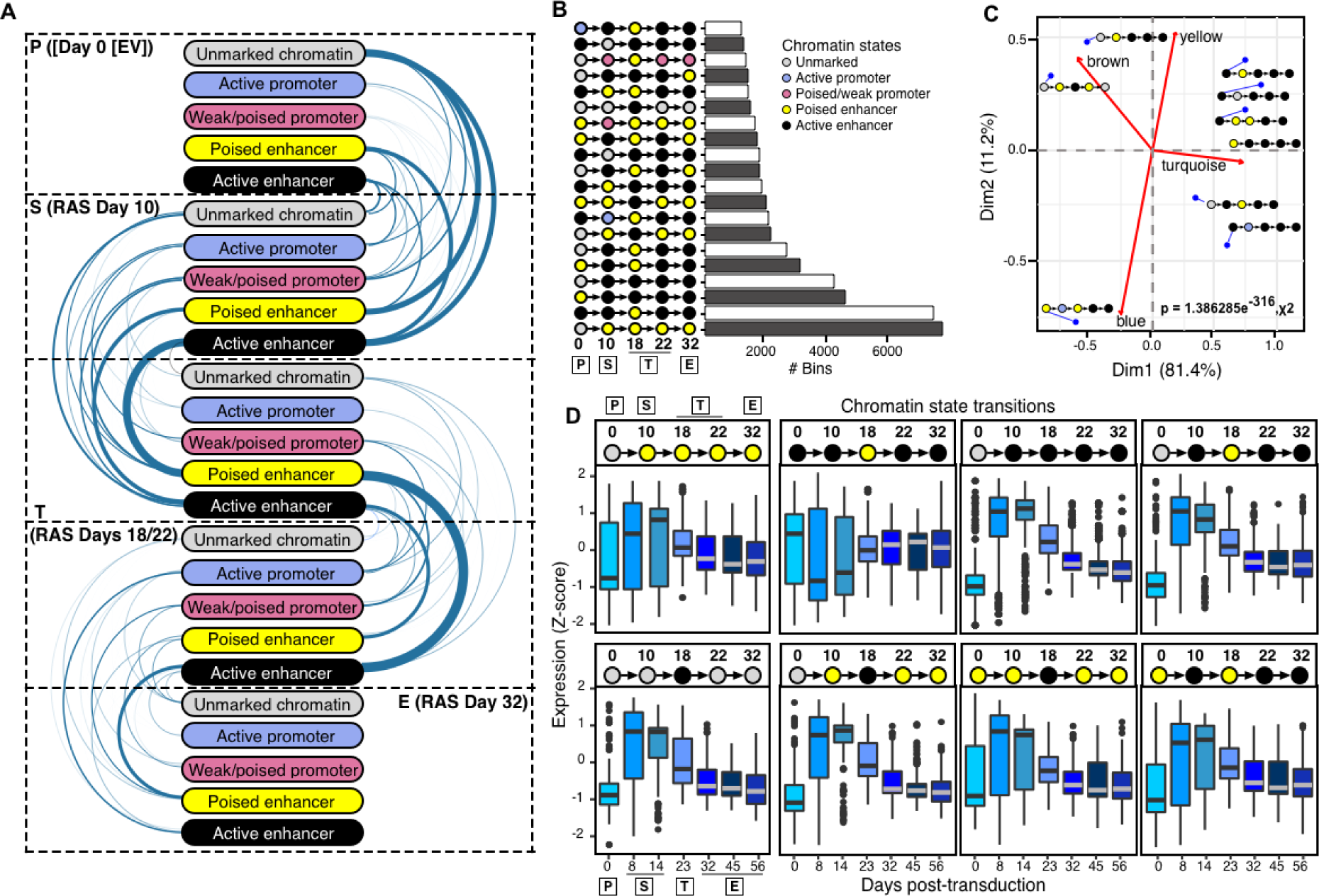
Enhancer remodeling dictates transcriptional transitions and OIS escape. **(A)** Arc-plot visualization of the chromatin state transitions at indicated stages as cells enter into and escape from OIS. The width of the edge is proportional to the number of 200 bp bins undergoing a given chromatin transition. **(B)** Histogram visualization of the number of 200 bp bins undergoing the top 20 most frequent chromatin state transitions during the escape from OIS. **(C)**, Asymmetric biplot of the CA between representative chromatin state transitions and gene expression modules. Nine representative and best projected (squared cosine > 0.5) chromatin state transitions are shown. Red lines arising from origin indicate the projections of gene expression modules (blue, brown, turquoise, yellow from Figure 1e). The statistical significance of the association was calculated using a chi-squared test. **(D)**, Integration of eight select chromatin state transitions (top pictograms; days 0, 10,18, 22 and 32 after H-RAS^G12V^ overexpression) with nearby gene expression output (row Z-score, boxplots; days 0, 8, 14, 23, 32, 45 and 56 after H-RAS^G12V^ overexpression). The centerline of the boxplots indicates the median, while the top and bottom edges correspond to the first and third quartiles. Whiskers extend from the box edges to 1.5x the interquartile range to the highest and lowest values. Chromatin state transition data (**A**-**D**) was computed from two independent immunoprecipitations per time-point per histone modification from pooled chromatin from 10 biologically independent experiments. Gene expression data (**D**) is the average of three biologically independent time series.

### ORGANIZED WAVES OF TRANSCRIPTION FACTOR ACTIVITY DEFINE ESCAPE FROM OIS

Transcription factors form dynamic hierarchical networks that finely control the duration and output of gene expression during cell fate transitions ^35–38^. This property of TF networks offers windows of opportunity to manipulate these transitions ^39–43^. To define the dynamics of TF networks during senescence escape, we first characterized the expression pattern of TFs in our time course. We identified 128 differentially expressed TFs in at least one stage as cells entered into and escaped from OIS (**Fig S4A)**. Our analysis revealed concordant upregulation of 45 TFs in the senescence-specific turquoise module, with concomitant downregulation of 83 proliferation-associated TFs in the blue, brown, and yellow modules during OIS. Expression of these TFs in cells that escaped from OIS stabilized to levels distinct from normal control cells, revealing that escape from OIS generates a transcriptome that allows normal somatic cells to proliferate, despite expression of oncogenic RAS (**Figure S4A**). Next, we integrated TF expression profiles and their differential binding activity at enhancers. We detected 339 TFs with differential activity throughout the OIS escape time series. While most of the binding dynamics were due to constitutively expressed TFs (289), only 50 differentially expressed TFs significantly contributed to this binding activity. These data suggest that cells leverage their existing TF networks to orchestrate the OIS response (**Figure 3A, Figure S4A**). In line with this interpretation, we observed organized waves of TF activity that defined each of the stages of OIS escape (i.e., P, S, T, and E). AP1 TFs, the OIS master regulators^27^, occupied the vast majority of enhancers and exhibited maximal binding activity during the senescence (S) and transition stages (T), then returning to near basal levels after OIS escape (**Figure 3A**). The E2F family of TFs exhibited a reciprocal behavior, being active only in proliferating control cells and after cells escaped from OIS, consistent with their role in promoting cell proliferation (**Figure 3A**). In contrast, POU and FOX family TFs exhibited peak activity shortly before cells escaped from OIS, indicating a potential role of these TFs in senescence escape (**Figure 3A**).

**Figure 3.**
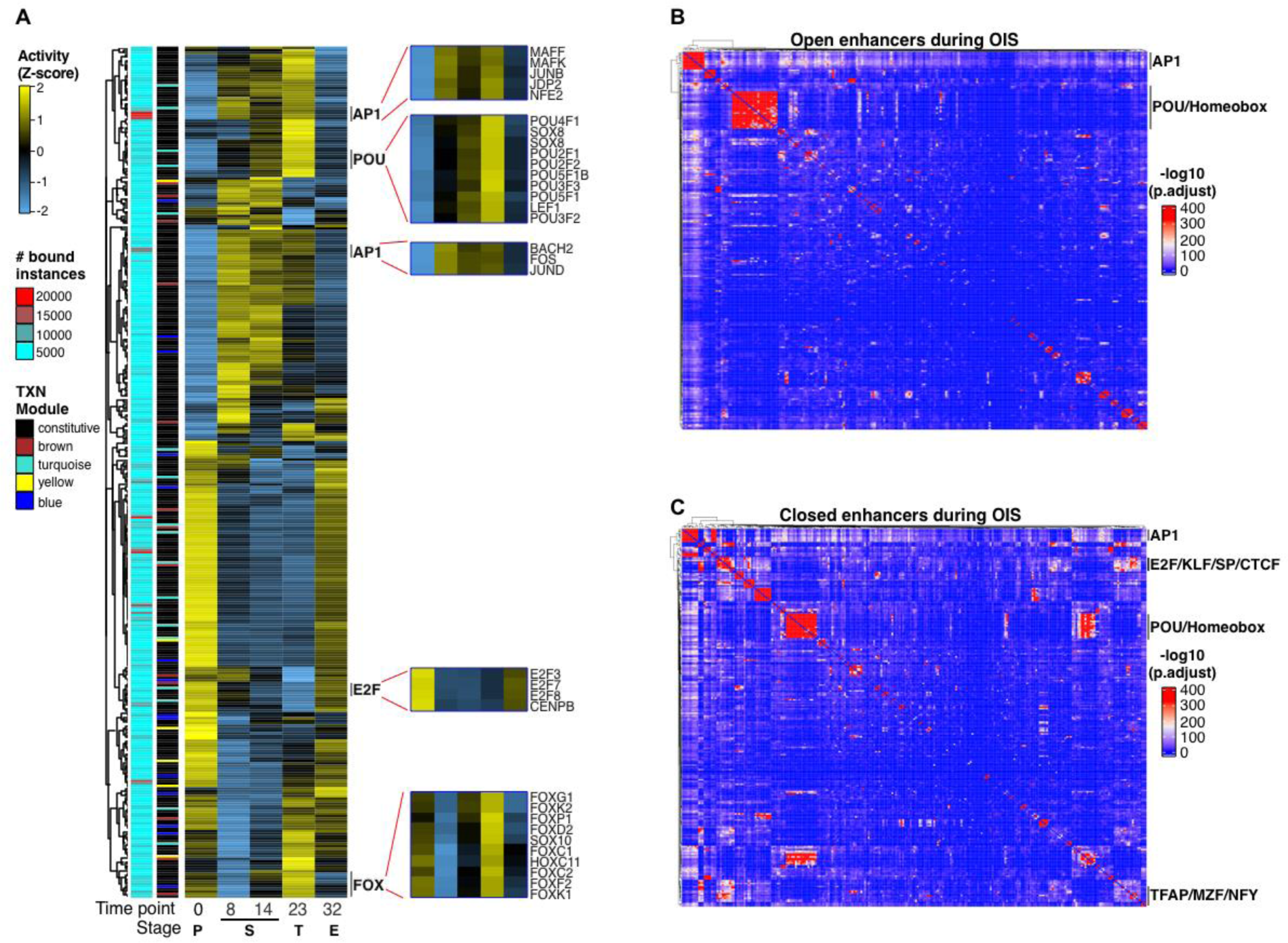
Organized waves of TF activity define OIS escape. **(A)** Heatmap showing the differential TF chromatin binding activity (row Z-score) at enhancers at each stage of OIS escape. Only expressed TFs were considered in the analysis. The annotations on the left show the number of bound instances per TF and their gene expression (TXN) category (i.e., constitutively expressed (black) or differentially regulated according to the module color code shown in Figure 1e). Insets show the chromatin binding activity of representative TFs families. **(B,C)** TF co-binding matrices at open **(B)** and closed **(C)** enhancers in cells entering and escaping from OIS. All binding instances across time points were collapsed onto the matrix and clustered using Ward’s aggregation criterion. The corresponding *q* values were projected onto the clustering and are color-coded based on significance calculated using a hypergeometric distribution test. TF footprinting **(A-C)** and differential chromatin binding activity (**a**) were performed on pooled ATAC-seq datasets from two biologically independent time-series experiments.

To gain insights into the TF network hierarchy, we collapsed all TF binding instances across time points and computed TF co-binding interactions in open and closed enhancers. Consistent with our previous findings ^27^, AP1 TFs mediated interactions with most other TFs in open and closed enhancers (**Figures 3B,C** and **Figures S4B,C**). In addition, we identified a distinct cluster involving AP1, POU, and other Homeobox-containing TFs in open and closed enhancers, suggesting that these TFs regulate enhancer dynamics. Interestingly, a cluster involving E2F, KLF, SP, TFAP, NFY, and CTCF TFs was apparent only in closed enhancers, indicating that they play a role in regulating cell cycle-associated genes independently of AP1 (**Figure 3C** and **Figure S4C**).

To visualize the hierarchical structure of the TF network during OIS escape, we constructed effector TF networks by focusing on the temporal co-binding interactions involving transcriptionally upregulated TFs from the OIS-specific gene module (**Figure S4A**, turquoise module), reasoning that some of these TFs likely play a role in escape from OIS. To this end, we used two different approaches focusing on: i) dynamically regulated enhancers of genes found in the transcriptional modules identified in Figure 1e (blue, brown, turquoise, and yellow modules) and ii) opening/closing enhancers at OIS irrespective of their relationship to the gene modules. Regardless of the approach, the structure of the TF networks was remarkably conserved, with AP1 TFs exclusively occupying the top layer and variable composition of the core and bottom layers (**Figures 4A-D, Figures 5A,B**). The organization and complexity of the networks, however, differed between transcriptional module- and enhancer-linked TF networks. Enhancer-linked TF networks occupied substantially more regions than their transcriptional module-coupled counterparts. Yet, they were simpler in their composition as AP1 interacted directly with TFs in the bottom layer or through 2 intermediary nodes (MZF1 and STAT3/STAT5 for open enhancers; IRF1/STAT1 and STAT3 for closed enhancers) in the core layer (**Figures S5A,B**). In line with our TF co-binding analysis, we confirmed the presence of a KLF/SP/CTCF/E2F subnetwork on closed enhancers that operated independently of AP1 (**Figure S5B**). TF regulation at enhancers of gene modules was more complex, with gene module-specific TF composition of the core layer (**Figures 4A-D**). Counterintuitively, we observed an inverse correlation between TF network complexity and gene module size. TF networks controlling the expression of the blue and turquoise modules (**Figures 4A,C**, 604, and 953 genes) had a simple architecture, with single-level core layers directly linking to the bottom layer TFs. In contrast, brown and yellow modules (**Figures 4B,D**, 310, and 247 genes) featured highly interconnected multilevel core layers interacting with the bottom layer TFs. In addition, a common feature of all networks was an overrepresentation of a subset of TFs (POU2F2, ZNF143, PRDM1, RBPJ, MTF1) in the core and bottom layers, suggesting a critical role for these TFs in mediating the escape from OIS.

**Figure 4.**
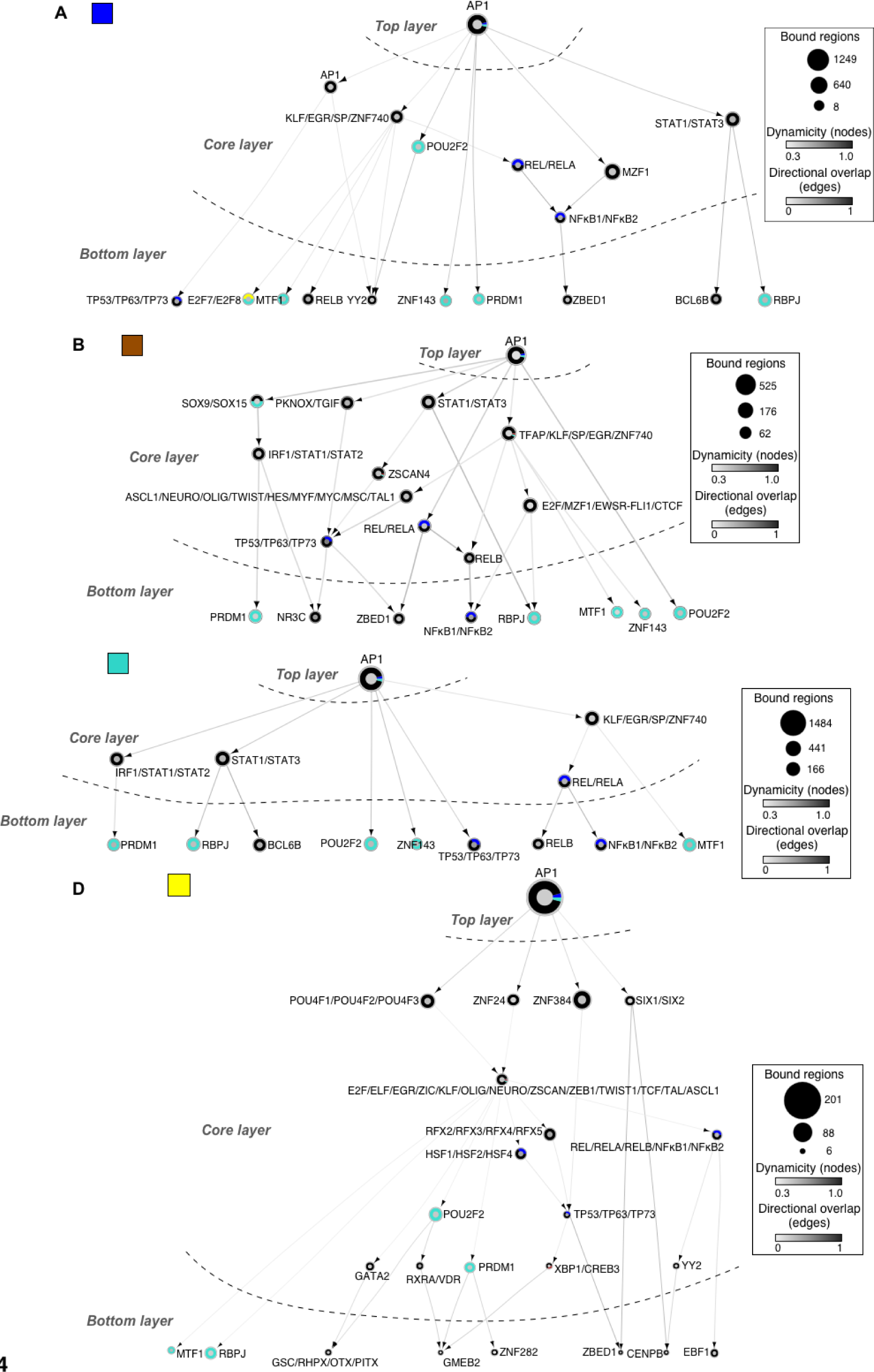
Hierarchical TF networks define transcriptional dynamics of OIS escape. (A-D) Effector TF networks for each gene expression module (Figure 1e). TFs (nodes) are represented as circles. Oriented edges (arrows) connecting nodes indicate that at least 15% of the regions bound by a given TF in the bottom and core layers were bound by the interacting TF in the core and top layers, respectively, at the same or previous time points. Strongly connected components (SCCs) are represented as a single node to facilitate visualization. The fill color of the node’s inner circle is based on the normalized dynamicity of TFs. The fill color of the outer ring indicates whether the TF is constitutively expressed (black) or belongs to a transcriptomic module (blue, brown, turquoise, and yellow). The node’s size is proportional to the bound regions by a given TF. Each network has three layers: i) top layer with no incoming edges, ii) core layer with incoming and outgoing edges, and iii) bottom layer with no outgoing edges. Networks were generated from pooled ATAC-seq data sets from two biologically independent time series.

**Figure 5.**
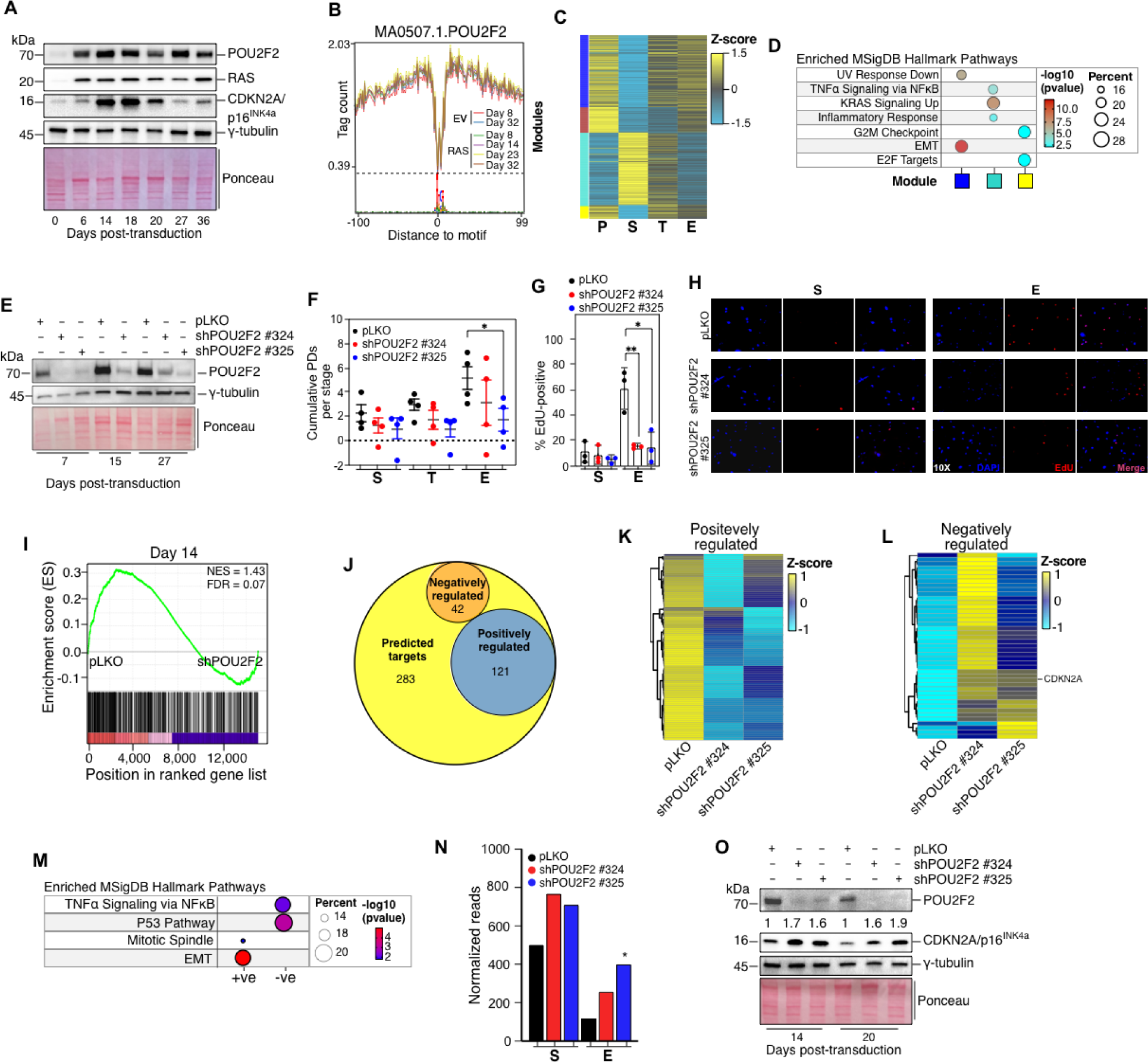
POU2F2 promotes OIS escape. **(A)** Western blot analysis of POU2F2, H-RAS^G12V^ and CDKN2A/p16^INK4a^ protein levels in GM21 fibroblasts undergoing OIS escape. γ-tubulin and Ponceau are shown as loading controls. One representative blot of three biologically independent experiments is shown. **(B)** Genome-wide POU2F2 enhancer-binding during OIS escape and time-matched controls. Footprinting was performed on pooled ATAC-seq data sets from two biologically independent experiments. **(C)** Expression heatmap of POU2F2 gene targets within the gene expression modules identified in Figure 1e. **(D)** Functional over- representation analysis map showing significant associations of the Molecular Signatures Database (MsigDB) hallmark gene sets with each POU2F2 gene target in the indicated modules. Circle fill is color-coded according to the FDR-corrected *p-value* from a hypergeometric distribution test. Circle size is proportional to the percentage of genes in each MsigDB gene set found within each gene module. **(E)** Western blot analysis of POU2F2 expression in RAS-expressing GM21 fibroblasts constitutively expressing non-targeting and POU2F2-targeting shRNAs. γ-tubulin and Ponceau are shown as loading controls. One representative blot of four biologically independent experiments is shown. **(F)** Cumulative population doubling plot at the OIS (S, days 8-16), transition (T, days 18-25), and escape (E, days 26-35) phases in GM21 fibroblasts constitutively expressing non-targeting and POU2F2- targeting shRNAs. Population doublings within each time interval per stage of four biologically independent experiments were included in the analysis. Statistical significance was calculated using a two-tailed unpaired Student’s *t*-test, comparing cells expressing POU2F2-targeting shRNAs to control shRNA-expressing cells at each stage (S, T, and E). *, *p* = 0.0412. **(G)** Histograms of EdU incorporation of GM21 fibroblasts constitutively expressing non-targeting and POU2F2-targeting shRNAs at the senescence (S) and OIS escape (E) stages. Statistical significance was calculated using a two-tailed unpaired Student’s *t*-test, comparing cells expressing POU2F2-targeting shRNAs to control shRNA-expressing cells at each stage (S and E) from three biologically independent experiments. **, *p* = 0.0088. *, *p* = 0.0173. **(H)** Representative immunofluorescence micrographs of EdU incorporation at the OIS and escape phases in GM21 fibroblasts constitutively expressing non-targeting and POU2F2-targeting shRNAs. **(I)** GSEA showing normalized enrichment score (NES) plots and FDR values for POU2F2 predicted target genes (**c**) in transcriptomes of pLKO and shPOU2F2-expressing cells at day 14 after RAS overexpression. Statistical evaluation of GSEA results was based on a nonparametric Kolmogorov–Smirnov test. **(J)** Intersections of positively and negatively regulated POU2F2 target genes within the predicted gene set defined in (**c**). **(K,L)** Expression heatmap of positively (k) and negatively (l) POU2F2-regulated genes. **(M)** Functional over- representation analysis map showing significant associations of the Molecular Signatures Database (MsigDB) hallmark gene sets with positively (+ve) and negatively (-ve) regulated POU2F2 target genes. **(N)** Histogram plot showing the normalized *CDKN2A* reads in cells as described in i. *, adjusted p value = 3.05 x 10^-5^, Wald test. The average of two biologically independent experiments is shown. **(O)** Western blot analysis of POU2F2 and CDKN2A/p16^INK4a^ expression in RAS-expressing GM21 fibroblasts constitutively expressing non-targeting and POU2F2-targeting shRNAs. γ-tubulin and Ponceau are shown as loading controls. Numbers on top of CDKN2A/p16^INK4a^ panel show densitometric fold increase relative to pLKO at each time point. One representative blot of three biologically independent experiments is shown.

In summary, our TF network analysis provides an information-rich yet simple representation of the complex relationships between TF binding dynamics and gene expression output as cells enter and escape from OIS, exposing TFs that may be decisive for OIS escape.

### POU2F2 PROMOTES ESCAPE FROM OIS

POU2F2, a member of the POU family of TFs ^44–46^, emerged as a potential candidate to promote OIS escape because it was transcriptionally upregulated shortly before cells escaped from OIS, was among the TFs with the highest differential binding activity at enhancers as cells transitioned to escape from OIS, and because of its direct connection to AP1 in the TF networks (**Figure S4A**, **Figure 3A, Figure 4,** and **Figure S5**). To corroborate the potential role of POU2F2 in OIS escape, we first used the Dynamic Regulatory Event Miner (DREM) ^47^ and Integrated Motif Activity Response (ISMARA) ^48^ software tools to predict transcriptional regulators from our time-series gene expression data in an unbiased fashion. Indeed, DREM identified POU2F2 as a transcriptionally induced TF that regulates cell cycle and cytoskeleton genes at the transition stage (days 14-23, **Figures S6A,B**). Further, ISMARA predicted peak POU2F2 activity between days 23 (transition) and 32 (escape) of the time series (**Figure S6C**). Next, we measured POU2F2 protein expression levels by immunoblotting. POU2F2 was barely detectable in control cells but increased dramatically as cells entered, remained in, and escaped from OIS (**Figure 5A**). The protein level of the senescence arrest gatekeeper CDKN2A (alias p16^INK4a^) was upregulated only in the senescence phase, declining to levels observed in control cells as cells escaped from OIS, which is consistent with CDKN2A mRNA expression pattern (**Figure 5A**, **Figure S1J**). Digital TF-footprinting of ATAC-seq datasets, an approach that faithfully recapitulates ChIP-seq-based TF-binding profiles as demonstrated by us previously ^49^, revealed a time-dependent increase in the chromatin binding activity of POU2F2 in oncogenic RAS-expressing cells, peaking at day 23 (transition) (**Figure 5B**). POU2F2 binding was dynamic, associating with different regions and genes, including CDKN2A, throughout the various stages as cells entered and escaped from OIS (total of 446 genes, **Figures S6D,E**).

Annotation of POU2F2 footprints identified 168, 182, and 32 target genes in the blue, turquoise, and yellow modules, enriching pathways relevant to OIS escape, including ‘EMT,’ ‘Inflammatory Signaling’ and ‘E2F targets’ (**Figures 5C,D**).

To directly test whether POU2F2 controls OIS escape, we silenced its expression before induction of OIS using two independent shRNAs and, as controls, also using non-targeting shRNAs. The expression of POU2F2-targeting shRNAs did not affect the proliferation rate of normal control cells (**Figures S6F,G**). By contrast, POU2F2 depletion stabilized OIS as early as the transition stage by significantly delaying or completely blocking oncogenic RAS expressing cells from escaping OIS in a manner that was dependent on silencing efficiency (**Figures 5E-H**). To understand the mechanism by which POU2F2 promotes escape from OIS in greater detail, we performed RNA-seq on control and shPOU2F2-expressing cells at days 14 (S) and 27 after (E) after RAS induction. Gene set enrichment analysis (GSEA) demonstrated that POU2F2 regulated a subset of its predicted targets (163/446 [36.5%] at day 14; 198/446 [44.3%] at day 27), in line with differential regulation of target genes by TFs ^50^ (**Figures 5I,J** and **Figures S6H,I**). POU2F2 acted primarily as a transcriptional activator but also negatively regulated the expression of a smaller subset of genes at both time points (**Figures 5K,L** and **Figures S6J,K**). Functional overrepresentation analysis of the validated POU2F2 gene targets confirmed its role in mediating escape from OIS, as it promoted the expression of genes involved in the EMT and cell proliferation, while it downregulated genes involved in inflammatory signaling and the P53 pathway (**Figure 5M**). Within this latter class, we identified CDKN2A/p16^INK4a^ as a critical POU2F2 target gene whose mRNA and protein levels increased upon POU2F2 downregulation (**Figures. 5N,O**). Overall, our results reveal that POU2F2 plays a central role in promoting escape from OIS by regulating cell identity, SASP- and cell cycle- genes.

### POU2F2 PROMOTES AN INFLAMMATORY GENE EXPRESSION PROGRAM IN COLORECTAL CANCER

To gain further insights into the role of POU2F2 during oncogenic RAS-driven carcinogenesis, we analyzed clinical, transcriptional, and chromatin accessibility data of human colorectal cancers (CRC) from The Cancer Genome Atlas (TCGA). We focused on CRC, not only because *RAS* gene mutations are present in a subset of aggressive cancers ^51–53^, but also because animal models of *KRAS*-driven CRC develop neoplastic lesions comprised of senescent cells ^54^, therefore modeling a scenario during which senescent cells may escape from OIS *in vivo*.

Consistent with recent evidence implicating POU2F2 in facilitating cancer progression in various tissues, including the brain, stomach, and lung, ^55–57^, increased *POU2F2* mRNA expression in primary CRCs was associated with decreased overall survival (**Figure 6A**). To understand how POU2F2 contributes to CRC, we performed footprinting analysis on chromatin accessibility datasets of 12 normal human colon tissue (from 5 donors) and 38 human CRCs, which revealed significantly upregulated POU2F2 chromatin binding activity in CRCs relative to normal tissue (**Figure 6B**). Annotation of POU2F2 footprints and integration with transcriptional data from primary CRC tumors identified a proliferative and inflammatory expression program composed of 1,712 differentially expressed genes (**Figure S7A**). Of these, we identified a POU2F2 signature comprised of 110 genes that overlapped with the POU2F2 targets identified in our *in vitro* OIS escape model. This OIS escape-indicating gene signature was characterized by overexpression of inflammatory and EMT-associated genes (**Figures S7B,C**), which are known to promote CRC progression ^58–61^. Furthermore, GSEA using this POU2F2-driven OIS escape gene signature revealed activation of IL17 and WNT signaling pathways, also known as drivers of CRC development ^62–66^ (**Figure 6C**).

**Figure 6.**
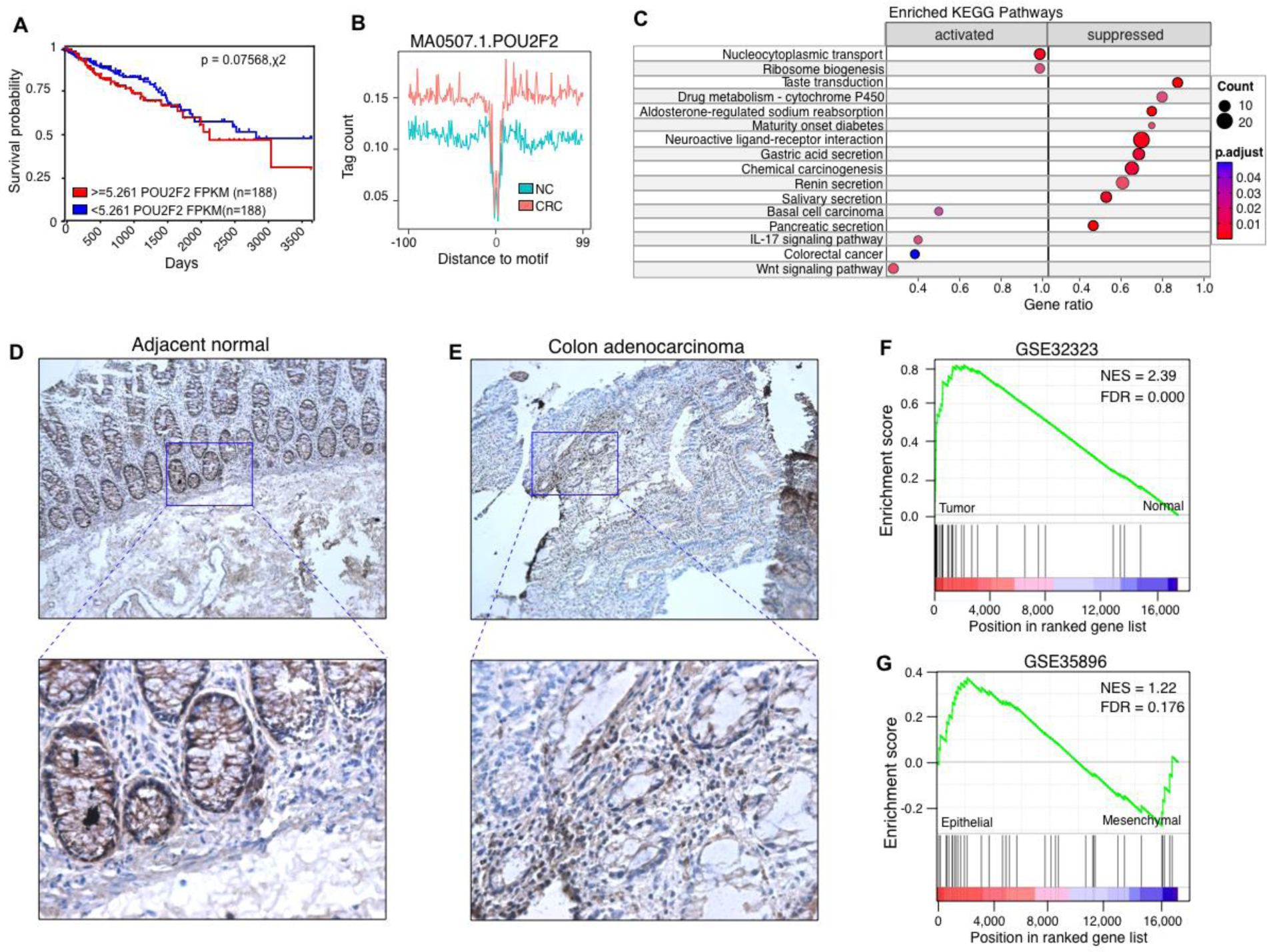
POU2F2 drives an inflammatory gene expression program in colorectal cancer. **(A)** Kaplan-Meier plot showing overall survival comparing colorectal cancer (CRC) patients with low (blue) and high (red) normalized *POU2F2* mRNA expression in primary tumors. Statistical significance was conducted using a χ^2^ test. *N* for each POU2F2 expression group is indicated in the plot. **(B)** Genome-wide meta profiles of POU2F2 binding at accessible chromatin in normal colon tissue (NC) and CRC. Footprinting was performed on 12 normal colons (5 donors) and 38 tumor DNAse-seq and ATAC-seq datasets. **(C)** Gene set enrichment analysis (GSEA) map showing significant associations of KEGG gene sets with a POU2F2 escape gene signature in CRC samples. Circle fill is color-coded according to the adjusted *p-value* based on a nonparametric Kolmogorov-Smirnov test. Circle size is proportional to the number of POU2F2 target genes within each KEGG gene set (rows). POU2F2 CRC gene targets were identified using gene expression data from 101 normal colon and 631 primary tumors deposited in the TCGA using the Xena Differential Gene Expression Analysis Pipeline. POU2F2 targets in CRC were then overlapped with POU2F2 targets in GM21 fibroblasts that escaped from OIS, and the union of these two sets was analyzed. **(D,E)** Representative micrographs at 10X (top) and 40X (bottom) magnifications of Hematoxylin and immunohistochemical analysis of POU2F2 expression in adjacent non-tumor biopsy and adenocarcinoma from a patient that underwent a colectomy procedure. **(F,G)** GSEA showing normalized enrichment score (NES) plots and FDR values for POU2F2 escape gene signature in transcriptomes (GSE32323) of CRC tumors (n=17) and normal tissue controls (n=17) profiled after colectomy procedures (**F**), and transcriptomes (GSE35896) of CRC tumors with mesenchymal (n=38) and epithelial (n=24) gene signatures (**G**). Statistical evaluation of GSEA results was based on a nonparametric Kolmogorov–Smirnov test.

To validate these findings, we analyzed POU2F2 levels by immunohistochemistry (IHC) in normal colon tissue, dysplastic polyps, high-grade colonic dysplasias, and colon adenocarcinoma from four different patients. We observed moderate cytoplasmic/membranous POU2F2 staining in enterocytes and endocrine cells of glands of the colonic mucosa, as well as moderate nuclear staining in sporadic mucosal lymphoid cells in non-tumor tissue (**Figures 6D,E** and **Figures S7D-H**), consistent with published data of POU2F2 distribution in normal human colon tissue ^67, 68^. In contrast, increasingly widespread, strong POU2F2 nuclear positivity was evident in dysplastic colonic tissue and particularly in colon (adeno)carcinoma (**Figures 6D,E,** and **Figures S7D-H**). Furthermore, using publicly available transcriptome datasets, we found that the POU2F2-driven OIS escape gene signature could separate tumors from normal colon tissue ^69^ and, to a lesser extent, previously defined CRC tumor subtypes with epithelial and mesenchymal character ^70^ (**Figures 6F,G**). In summary, integrative analysis of epigenomic and transcriptomic datasets from human CRCs and OIS escape highlight POU2F2 activation and its associated OIS-escape gene signature as prognostic biomarkers and potential parameters for CRC progression in humans.

### AN EPIGENETIC MEMORY DEFINES POST-SENESCENT CELLS AND IS DETECTABLE IN COLORECTAL CANCER

Although senescence escape may be a path to overt carcinogenesis ^3, 18^, direct evidence of senescence escape as a potential cause of cancer progression in humans remains elusive. The reproducibility of the gene expression trajectories and the persistence of enhancer states after OIS escape prompted us to evaluate whether an epigenetic memory of this event could be identified in OIS escapee cells. To this end, we interrogated our chromatin accessibility datasets, focusing on enhancers. The differential analysis identified two modules comprising 27,117 enhancers whose accessibility changed during OIS (**Figure 7A**). Visualization of differentially accessible chromatin trajectories by PCA revealed virtually identical transitions to those identified in gene expression datasets of cells that entered and escaped OIS. In contrast, time-matched control cells remained stable (**Figure S8A**). Interestingly, integrating chromatin accessibility dynamics with transcriptional modules revealed contributions of both opening and closing enhancers to the gene expression output, confirming the role of local chromatin state dynamics in determining gene expression levels ^71^ (**Figure S8B**). We next performed time-point pair-wise differential analysis to identify enhancer regions that permanently change accessibility as cells enter and escape from OIS. This analysis identified 1,926 differentially accessible regions in OIS that were also preserved in cells that escaped from OIS, which we termed senescence-associated chromatin scars (SACS) (**Figures 7B,C**, **Figures S8C-F**). Most of these SACS localized to genomic regions poised to become active enhancers (**Figure 7D**) and enriched for senescence- and cancer-associated pathways in functional enrichment analyses using two independent databases (Molecular Signatures ^72^ and KEGG pathways ^73^) (**Figures 7E,F**). Representative genome browser snapshots of opening and closing SACS are shown (**Figures S8G,H**). These results demonstrate that OIS escapee cells store an ‘epigenetic memory’ of OIS.

**Figure 7.**
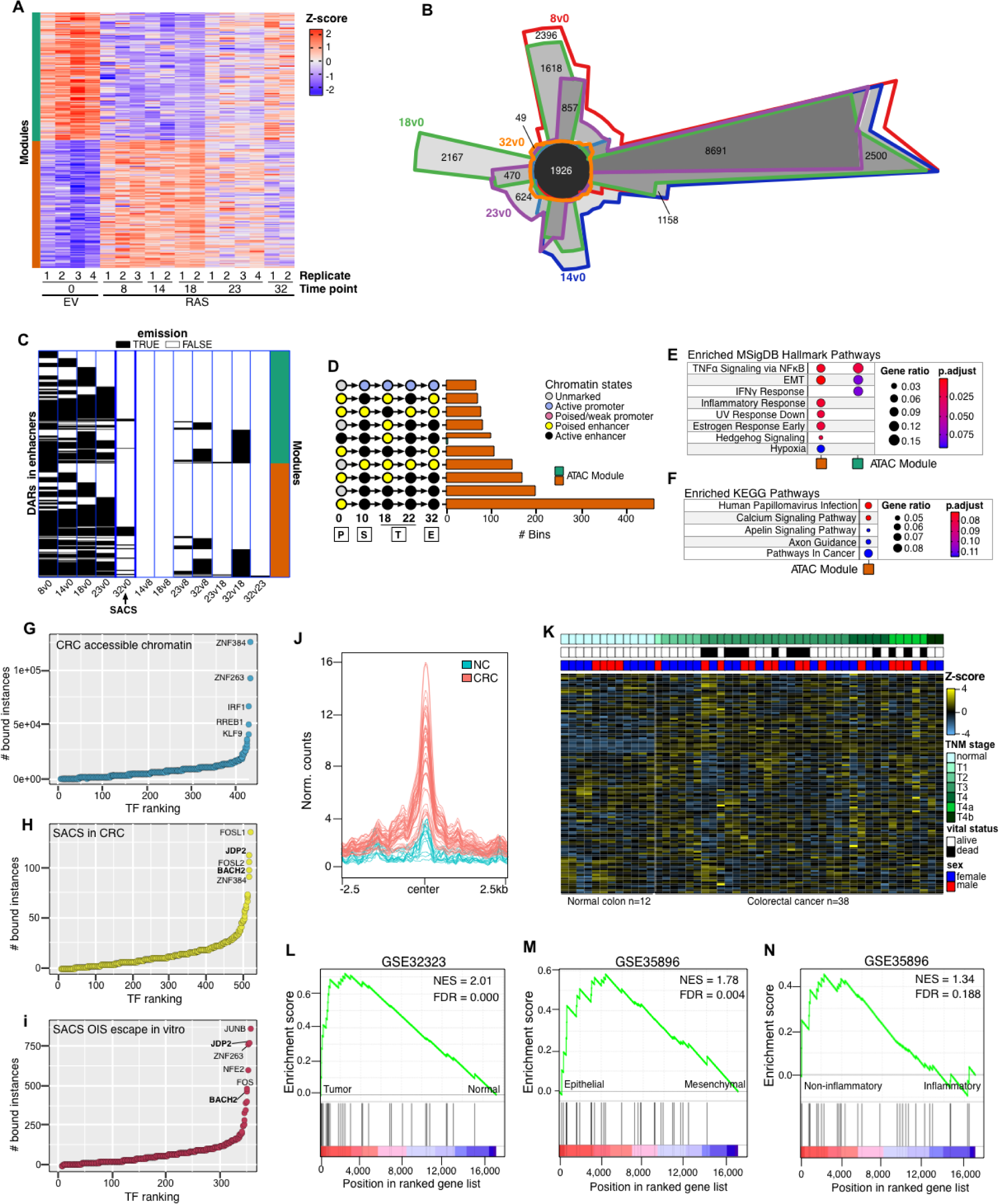
SACS define post-senescent cells and are present in colorectal cancer. **(A)** Heatmap showing the chromatin accessibility modules at enhancers per time point as cells enter into and escape from OIS. 27,117 differentially accessible regions were identified. Biologically independent replicates are numbered. **(B)** Chow-Ruskey diagram showing intersections and disjunctive unions of differentially accessible chromatin regions at each time point measured as cells enter into and escape from OIS, relative to the empty vector control (day 0). The black center circle depicts 1,926 senescence scars. **(C)** True/False emission heatmap visualization of differentially accessible genomic regions at each chromatin accessibility module at enhancers for each time point pairwise comparison, as indicated at the bottom of the heatmap. A true emission means a region is differentially accessible relative to the indicated reference for each chromatin accessibility module. SACSs span horizontally for all time point comparisons relative to day 0 (five leftmost lanes of the heatmap). **(D)** Histogram visualization of the number of 200 bp windows within senescence scars undergoing the top 10 most frequent chromatin state transitions. Functional over-representation analysis map showing significant associations of the Molecular Signatures Database (MSigDB) Hallmark **(E)** and Kyoto Encyclopedia of Genes and Genomes (KEGG) **(F)** gene sets with genes associated with SACSs in each chromatin accessibility module. Circle fill is color-coded according to the FDR-corrected p-value from a hypergeometric distribution test. Circle size is proportional to the ratio of genes in the database sets (MSigDB or KEGG) found within the genes associated with SACSs in each chromatin accessibility module. Data shown represent the number of biologically independent experiments per time point indicated in **(A)**. **(G-I)** Rank plots showing the summed binding instances of TFs in accessible chromatin, SACS in CRC, and SACS in OIS escapee cells. **(J)** Normalized metaprofiles of SACS in individual normal colon and CRC samples. **(K)** Annotated heatmap showing signal intensity (row Z-score) of the 114 CRC-specific SACS across individual samples and their association to AJCC TNM stage, vital status, and sex of the patient. *N* for normal colon and tumors is shown. **(L-N)** GSEA showing normalized enrichment score (NES) plots and FDR values for SACS-associated gene signature in transcriptomes (GSE32323) of CRC tumor (n=17) and their normal tissue controls (n=17) profiled after colectomy procedures (**l**), transcriptomes (GSE35896) of mesenchymal- (n=38) and epithelial-like (n=24) CRC tumors (**m**) and transcriptomes (GSE35896) of non-inflammatory (n=18) and inflammatory (n=5) epithelial-like CRC tumors (**n**). Statistical evaluation of GSEA results was based on the nonparametric Kolmogorov–Smirnov test. SACS-associated genes in CRC were identified using gene expression data from 101 normal colon and 631 primary tumors deposited in the TCGA using the Xena Differential Gene Expression Analysis Pipeline.

To test whether SACS are detectable in cancer, we sought to identify them in CRC-accessible chromatin. To this end, we performed differential chromatin accessibility analysis in normal colon and CRC, identifying six distinct modules (**Figure S8I**). Overlapping the genomic coordinates of SACS with accessible chromatin modules from normal colon and CRCs revealed an overlap of 114 SACS exclusively in CRC accessible chromatin modules (black, green, and cyan; **Figure S8J**). Importantly, relative to the overall TF binding distribution across the CRC accessible chromatin landscape, AP1 TFs were preferentially bound to SACS in CRC, consistent with their binding profile to SACS identified *in vitro* (**Figures 7G-I**). Furthermore, the signal distribution across SACS revealed increased intensity in CRC across all cancer stages relative to the normal colon (**Figures 7J,K**), indicating that SACS may originate in premalignant lesions. However, individual differences in the genomic region and tissue/tumor sample could be observed (**Figure 7K** and **Figure 8K**), likely due to the cellular heterogeneity of colonic tissue ^74^. To determine whether SACS in CRC could provide diagnostic information regarding disease state, we integrated CRC SACS with transcriptomic data from primary CRCs and identified a signature of 47 differentially regulated genes that we evaluated for its prognostic power as a biomarker in publicly available CRC datasets ^69, 70, 75^. Indeed, this SACS gene signature readily separated tumors from normal colon tissue ^69, 70, 75^. Furthermore, this gene signature distinguished epithelial subtype CRCs with good prognosis from mesenchymal subtype CRCs with poor prognosis and, to a lesser extent, non-inflammatory from inflammatory epithelial subtype CRCs (**Figures 7L-N**). Our integrative analysis therefore provides strong evidence for epigenomic features of senescence escape within the CRC epigenome with diagnostic and prognostic value.

## Discussion

Recent studies revealed that cellular senescence is not a stable barrier to cancer progression ^21, 24, 26, 52, 75–78^. However, the gene-regulatory mechanisms that control TF networks and promote escape from cellular senescence and whether post-senescent cells are present in human cancers are largely unknown. This knowledge, however, would prove highly useful not only for diagnostic and prognostic purposes in a variety of cancers that have been shown to develop features of OIS, but also for the development of therapies to modulate senescence-associated cell fate transitions during early cancer development and to target OIS escapee cells at later cancer stages. Our study reveals AP1 and POU2F2 as critical regulators of the transcriptional program that allow cells to escape from OIS and thus provides the foundation for understanding the processes that enable oncogene expressing cells to escape the barriers that constrain their continued proliferation.

A major finding of our study is that entry into and exit from OIS is precoded and mediated at the epigenetic level by AP1 pioneer TFs. This extends the role of AP1 TFs as master regulators of the senescence program ^27^ to critical mediators of senescence-associated cell fate transitions. We confirm that AP1 bookmarks prospective senescence enhancers, suggesting that it not only orchestrates entry into senescence but that it also promotes OIS escape by facilitating interactions of senescence-induced TFs with enhancer chromatin. These interactions, newly established during the senescence stage, likely execute the transcriptional programs required for senescence escape. This highly complex and orchestrated process follows reproducible transcriptional and epigenome trajectories, indicating that the capacity of cells to engage and disengage the senescent state is preestablished by TF-enhancer interactions throughout the epigenome. Thus, OIS and its associated cell fate transitions are not simply stress responses but they are essentially indistinguishable from other epigenetically precoded processes such as cell differentiation programs^79^.

Our data also reveal that escape from OIS is accompanied by a gradual downregulation of SASP-associated genes and re-expression of genes that promote cell cycle progression. SASP gene expression never returned to basal levels in cells that escaped from OIS but remained elevated at significantly higher levels than in normal cells. Therefore, cells that had escaped from OIS retained a secretory phenotype known to drive cancer initiation, proliferation, progression, metastasis, and resistance to therapy^22, 80^. As SASP factors have also been shown to induce stemness and cellular reprogramming through paracrine mechanisms ^15, 16^, our results raise the possibility that OIS-escaped cells, using their remnant SASP, render cells within their microenvironment more susceptible to further transformation by increasing their plasticity.

Our time-resolved integrative profiling revealed novel transcriptional regulators relevant to cancer development and epigenomic features with prognostic potential. Foremost, we identified POU2F2 as a critical factor that promotes OIS escape *in vitro* and provided compelling evidence that POU2F2 performs a similar function during CRC development, where its overexpression and increased activity are associated with a proliferative and inflammatory transcriptional program as well as decreased overall survival. Furthermore, this putative function of POU2F2 in CRC is consistent with recent evidence demonstrating its role in the progression of cancers in several tissues, including in stomach, lung, brain, and germinal B-cell lymphomas, where it regulates transcriptional networks that alter metabolic pathways, promotes cell proliferation and metastasis and activates cell differentiation programs ^55–57, 81^. Collectively, these findings highlight POU2F2 as a potential therapeutic target in CRC.

Another important finding is the identification of 114 AP1-associated epigenetic remnants of OIS in human CRCs, termed senescence-associated chromatin scars (SACS). Significantly, the discovery of SACS in CRCs suggests that at least a subset of the CRC samples analyzed in this study underwent a transient period of senescence during cancer development. Given AP1 TFs as master mediators of the senescence program^27, 82^, the preferential binding of AP1 TFs at SACS relative to the overall CRC accessible chromatin landscape supports this conclusion.

Furthermore, the CRC SACS gene signature distinguished patients with epithelial-type CRC from those with mesenchymal-type CRC and consequently predicted disease-free survival after surgery based on published data ^70, 83^. These findings, thus, underscore the prognostic and diagnostic power of senescence-associated gene signatures in CRC and support the beneficial aspects of senescence in the outcome of anticancer therapy ^6, 27, 84^. Future work will have to establish the generalizability of senescence-associated gene signatures as diagnostic and prognostic tools during disease progression and their ability to inform and direct therapeutic decision-making.

In summary, our work provides a comprehensive framework of the gene regulatory mechanisms governing OIS escape and its associated cell fate transitions in the context of cancer development. It also highlights the strength of integrative time-resolved profiling as an effective approach to improving (pre-)cancer diagnostics and prognostication and revealing therapeutic liabilities to modulate the senescence program for clinical benefit.

## Acknowledgments

R.I.M.-Z. is a Mexican National Researchers System (SNI) fellow. This study was supported by the National Cancer Institute of the NIH (R01CA136533). The content is solely the responsibility of the authors and does not necessarily represent the official views of the National Institutes of Health.

## Author contributions

R.I.M.-Z and A.S generated cell culture systems, performed ChIP-seq, ATAC-seq, and RNA interference experiments, and analyzed data. R.I.M.-Z. performed computational analyses on the microarray, ChIP-seq, ATAC-seq, ENCODE, and TCGA data, generated and prepared figures. P.-F.R processed ATAC-seq data and designed computational pipelines to analyze microarray and ChIP-seq data. A.S performed colony formation and telomerase activity experiments and analyzed data. A.S. and T.V performed EdU incorporation and TIF experiments and analyzed data. M.S. performed immunohistochemistry experiments. G.D. generated the microarray data. J.A.N generated the TF networks and analyzed ATAC-seq data. R.C. conducted surgeries on cancer patients, R.C. and M.G. provided tumor tissue and consulted with pathologic diagnoses.

## STAR Methods

### CELL CULTURE

Human somatic GM21 foreskin fibroblasts (Coriell Institute, Camden, NJ) and WI-38 human fetal lung fibroblasts (European Collection of Authenticated Cell Cultures, Porton Down, UK) were cultured in a DMEM medium containing 10% fetal bovine serum (FBS) and 1× penicillin/streptomycin (Corning) at 37 °C in a 2% oxygen atmosphere. GM21- and WI38-RAS^V12^ fibroblasts and their empty vector counterparts were generated by retroviral transduction as previously described in ^24^. Samples were collected and processed at the time points indicated in the main text. GM21 fibroblasts constitutively expressing non-targeting and POU2F2-targeting shRNAs (pLKO.1-Neo-CMV-tGFP, pLKO.1-Neo-CMV-tGFP-TRCN0000245324 and pLKO.1-Neo-CMV-tGFP-TRCN0000245325) (Millipore-Sigma, Saint Louis, MO) were generated by lentiviral transduction and selected with 400 µg/ml neomycin for seven days. Cells were subsequently retrovirally transduced with RAS^V12^ or empty vector previously described in ^24^.

### RNA AND MICROARRAYS

RNA from each time-point from GM21 fibroblasts undergoing escape from OIS was purified using the Macherey-Nagel Nucleospin RNA Plus XS kit according to the manufacturer’s instructions (Macherey-Nagel Inc, Allentown, PA). According to the manufacturer’s instructions, 100 ng RNA per sample was analyzed using Affymetrix Human Transcriptome Arrays 2.0 (Applied Biosystems, Santa Clara, CA).

### MICROARRAY DATA PREPROCESSING

Data were processed as described in ^27, 85^. Briefly, all microarrays were normalized simultaneously using the robust multi-array normalization algorithm implemented in the *oligo* package. Internal control probes were removed, and deciles of average expression were defined for the entire time series. Probes falling in the lowest four deciles of expression were removed. Identification and removal of unidentified sources of variability were performed using the *removebatcheffect* option of the limma package. Replicate similarity was evaluated at each preprocessing step using principal component analysis, bi-clustering using Pearson’s correlation with Ward’s aggregation criterion.

### SELF-ORGANIZING MAPS (SOMs)

SOM expression portraits were generated using the unsupervised machine learning method deployed in oposSOM^33^. Metagenes were visualized in a 60 x 60 grid of rectangular topology, wherein expression portraits are projected by metagene distance matrix similarity using a logarithmic fold-change scale. To verify that expressions portraits from GM21 fibroblasts escaping OIS differ from time-matched controls, we implemented the D-clustering feature of oposSOM, which reveals clusters of differentially expressed genes based on SOM units with local maxima relative to the mean Euclidean distance to their neighbors.

### DIFFERENTIAL GENE EXPRESSION ANALYSIS AND GENE CO-EXPRESSION NETWORKS

Normalized and preprocessed microarray expression data were analyzed for differential expression using limma. Due to the similarity of consecutively measured time points on PCA plots and SOM portraits, we collapsed them into four stages: proliferation, senescence, transition, and escape. We performed contrasts comparing the senescence, transition, and escape stages to the proliferation reference to identify differentially expressed genes. The statistical approach involves empirical Bayes-moderated t-statistics applied to all contrasts for each probe, followed by moderated F-statistic to test whether all the contrasts are zero, evaluating the significance of the expression changes observed ^86^. For significant probes, p-values were corrected for multiple testing with the FDR approach using a significance level of 0.005 as a cut-off. Probes matching these criteria that also exhibited a minimal absolute fold change of 1 were considered differentially expressed genes (DEGs). 2,118 DEGs were used as input for unsupervised clustering using WGCNA ^87^ using the ‘signed’ option with default parameters except for the soft-thresholding power, which was set to 19. The identified co-expressed gene modules were visualized by heatmaps, network, and PCA plots and functionally characterized using clusterProfiler ^88^ using the Molecular Signatures Database Hallmark and KEGG gene sets ^72, 73^.

### DREM AND ISMARA

Time-series microarray gene expression data were analyzed using DREM ^47^ and ISMARA ^48^. DREM was run using default settings except for the change thresholding parameter, set to the maximum-minimum setting with a minimum absolute expression change of 1. Gene Ontology analysis was performed with default settings with statistical significance calculated using randomization with 500 random samples. For ISMARA, raw CEL files were uploaded to the ISMARA server (https://ismara.unibas.ch/mara/) and ran using default parameters.

### HISTONE MODIFICATION CHIP-SEQ

Chromatin was collected from GM21 fibroblasts retrovirally transduced with an empty vector (referred to as day 0) or H-RAS^G12V^ at days 10, 18, 22, and 32 after transduction. A total of 0.8- 1 × 10^7^ cells (per time point, from 10 pooled biological replicates) were fixed in 1% formaldehyde for 15 min, quenched in 2 M glycine for an additional 5 min, and pelleted by centrifugation at 2,000 r.p.m., 4 °C for 4 min. Nuclei were extracted in extraction buffer 2 (0.25 M sucrose, 10 mM Tris-HCl pH 8.0, 10 mM MgCl2, 1% Triton X-100 and proteinase inhibitor cocktail) on ice for 10 min, followed by centrifugation at 3,000 × g at 4 °C for 10 min. The supernatant was removed, and nuclei were resuspended in nuclei lysis buffer (50 mM Tris-HCl pH 8.0, 10 mM EDTA, 1% SDS, and proteinase inhibitor cocktail). Sonication was performed using a Branson sonicator until the desired average fragment size (100–300 bp) was obtained. Soluble chromatin was obtained by centrifugation at 12,000 r.p.m. for 15 min at 4 °C, and chromatin was diluted tenfold. Immunoprecipitation was performed overnight at 4 °C with rotation using 1 × 10^6^ cell equivalents per immunoprecipitation using antibodies (5 µg) against H3K4me1 (Active Motif, Carlsbad, MA; Cat no. 39297) and H3K27ac (Active Motif, Carlsbad, MA; Cat no. 39133). Subsequently, 30 µl of Ultralink Resin (Thermo Fisher Scientific, Waltham, MA) was added and allowed to tumble for 4 h at 4 °C. The resin was pelleted by centrifugation and washed three times in low-salt buffer (150 mM NaCl, 0.1% SDS, 1% Triton X-100, 20 mM EDTA, 20 mM Tris-HCl pH 8.0), one time in high-salt buffer (500 mM NaCl, 0.1% SDS, 1% Triton X-100, 20 mM EDTA, 20 mM Tris-HCl pH 8.0), two times in lithium chloride buffer (250 mM LiCl, 1% IGEPAL CA-630, 15 sodium deoxycholate, 1 mM EDTA, 10 mM Tris-HCl pH 8.0) and two times in TE buffer (10 mM Tris-HCl, 1 mM EDTA). Washed beads were resuspended in elution buffer (10 mM Tris-Cl pH 8.0, 5 mM EDTA, 300 mM NaCl, 0.5% SDS) treated with RNAse H (30 min, 37 °C) and Proteinase K (2 h, 37 °C), 1 µl glycogen (20 mg/ml; Ambion, Austin, TX) was added and decrosslinked overnight at 65 °C. For histone modifications, DNA was recovered by mixing the decrosslinked supernatant with 2.2× SPRI beads followed by 4 min of incubation at room temperature. The SPRI beads were washed twice in 80% ethanol, allowed to dry, and DNA was eluted in 35 µl of 10 mM Tris-Cl pH 8.0. Libraries were constructed using an Accel-NGS 2S Plus DNA Library kit (21024; Swift Biosciences, Ann Arbor, MI) and amplified for 9 cycles. Libraries were then resuspended in 20 µl of low EDTA–TE buffer. Libraries were quality controlled in an Agilent Technologies 2100 Bioanalyzer (Applied Biosystems, Santa Clara, CA) and quantified using an Invitrogen Qubit DS DNA HS Assay kit (Q32854) (Thermo Fischer Scientific, Waltham, MA). Libraries were sequenced using an Illumina High-Seq 2500. Typically, 30–50 million reads were required for downstream analyses.

### ATAC-SEQ

The transposition reaction and library construction were performed as previously described in ^89^. Briefly, 50,000 cells from each time point of the OIS escape time course (two biological replicates) were collected, washed in 1× in PBS, and centrifuged at 500 × g at 4 °C for 5 min.

Nuclei were extracted by incubating cells in nuclear extraction buffer (containing 10 mM Tris-HCl, pH 7.4, 10 mM NaCl, 3 mM MgCl2, 0.1% IGEPAL CA-630) and immediately centrifuging at 500 × g at 4 °C for 5 min. The supernatant was carefully removed by pipetting, and the transposition was performed by resuspending nuclei in 50 µl of Transposition Mix containing 1× TD Buffer (Illumina, San Diego, CA) and 2.5 µl Tn5 (Illumina) for 30 min at 37 °C. DNA was extracted using a Qiagen MinElute kit (Qiagen, Germantown, MD). Libraries were produced by PCR amplification (12 cycles) of tagmented DNA using an NEB Next High-Fidelity 2× PCR Master Mix (New England Biolabs, Ipswich, MA). Library quality was assessed using an Agilent Bioanalyzer 2100 (Applied Biosystems, Santa Clara, CA). Paired-end sequencing was performed in an Illumina Hiseq 2500 system. Typically, 30–50 million reads per library were required for downstream analyses.

### RNA-SEQ

RNA from control and shPOU2F2-expressing GM21 fibroblasts at the senescence (day 14) and escape (day 27) phases was isolated with a Macherey-Nagel Nucleospin RNA Plus XS kit according to the manufacturer’s instructions (Macherey-Nagel Inc, Allentown, PA). The RNA integrity number (RIN) of all samples was at least 9. Libraries were generated using the NEBNext Ultra II Library Prep Kit (New England Biolabs, Ipswich, MA) and sequenced in an Illumina Hiseq 2500 system. 40 million reads per library were required for downstream analyses.

### PREPROCESSING OF RNA-SEQ, ATAC-SEQ AND CHIP-SEQ

Paired-end (RNA-seq and ATAC-seq) and single-end reads (ChIP-seq) were aligned to the hg19 version of the human genome using bowtie2 ^90^ using the local mode. Low-quality reads and adapters were removed using fastq-mcf v.1.0.5. Alignments were further processed using samtools v.1.1.1, and PCR and optical duplicates were removed with PicardTools v.2.2.2.

Enriched regions for histone modifications and ATAC-seq were identified using macs v.2.2.7.1 (macs2 callpeak --nomodel --shiftsize --shift-control--gsize hs -p 1e-3). The identified peaks were subsequently processed using the irreproducibility discovery rate (IDR) pipeline ^91^, generating time point-specific reproducible peak sets for histone modifications and ATAC-seq at each time point. For RNA-seq samples, reads were counted using *summarized overlaps* and normalized with DESeq2 for visualization.

### CHROMATIN STATE TRANSITIONS

To identify and quantify combinatorial chromatin state dynamics, we analyzed the genome-wide combinations of H3K4me1, H3K27ac, and ATAC-seq signals on IDR-controlled reproducible peaks of GM21 escaping OIS using the chromstaR package, which relies on implementing a univariate Hidden Markov Model (HMM) with two hidden states (unmodified and modified) ^92^.

Briefly, the algorithm was run on differential mode and conFigd to partition the genome in 200 bp non-overlapping bins and count the number of reads of histone modification/ATAC-seq mapping into each bin at the time points indicated in the main text, and modeled using a univariate HMM based on a two-component mixture, the zero-inflated binomial distribution.

Subsequently, a multivariate HMM assigns every bin in the genome to one of the multivariate _components considering 2_(5-time points x 3 genomic enrichment variables [histone modifications and ATAC-seq]) _possible_ states. To verify the chromatin states linked to histone modifications/ATAC-seq in GM21 fibroblasts undergoing escape from OIS, we performed extensive comparisons with chromatin state data generated in NHLF cells from the ENCODE project. To evaluate the relationship between chromatin state dynamics and transcriptional output, we first identified nearby genes to the select chromatin state transitions shown in Figs. 2c-d, further selected genes belonging to the identified co-expression modules, and visualized their expression dynamics relative to chromatin state transitions using correspondence analysis (CA, Figure 2c) and boxplots (Figure 2d). For CA, results were visualized on an asymmetric biplot using FactoMineR after filtering for the top contributing chromatin state transitions (square cosine >0.5).

### DIFFERENTIAL CHROMATIN ACCESSIBILITY, SENESCENCE SCARS, AND HIERARCHICAL CLUSTERING

To highlight peaks with differential accessibility (DA) over time, we used DESeq2 ^93^. ‘Senescence scars’ were defined as the intersection of DA peaks when comparing each time point in the dataset (Days 8, 14, 18, 23, and 32) with peaks identified at Day 0. Venn diagram plots were generated by the R package Vennerable. To identify modules of peaks with correlated accessibility profiles over time, we used WGCNA. We set the ‘minimum cluster size’ and the ‘deepSplit’ parameters to 130 and 3. We obtained the value of the ‘soft threshold’ parameter, set at 20, using the elbow method recommended by the developers. By visually inspecting the obtained modules, we defined the ‘dissimilarity threshold’ parameter as 0.20.

### ATAC-SEQ ANNOTATION AND TRANSCRIPTOME INTEGRATION

We determined the closest gene to each DA peak using the R package ChIPseeker ^94^. We used the clusterProfiler R package to perform an over-representation analysis of each identified DA peak module based on the Molecular Signatures Database. We integrated our DNA accessibility and transcriptomic datasets by inspecting the overlap of DA peaks target genes with identified DE genes and generated a river plot using the R package NetworkD3 to observe the association between the dynamic profiles obtained by the hierarchical clustering of each dataset.

### NORMALIZED GENOME BROWSER TRACKS

For genome track visualizations, RNA-seq, ChIP-seq and ATAC-seq alignments were normalized using deeptools v3.3.1 ^95^ using the RPGC approach to obtain 1X coverage (bamCoverage -b –normalizeUsing RPGC –effectiveGenomeSize 2864785220 – ignoreDuplicates –binSize 10 –verbose -o).

### TRANSCRIPTION FACTOR FOOTPRINTING AND DIFFERENTIAL BINDING ACTIVITY

Footprinting, TF motif enrichment, and differential binding activity were performed with HINT-ATAC ^96^ using the JASPAR position weight matrix database for vertebrate TFs ^97^ on merged ATAC-seq datasets (at least two biological replicates) of the indicated time points of GM21 fibroblasts undergoing escape from OIS. HINT-ATAC uses position dependency models (PDM) to estimate the bias of Tn5 cleavage events and incorporates nucleosomal density and strand-specific information to increase footprinting quality. We limited our footprinting and differential binding activity analyses to enhancer regions undergoing chromatin state transitions defined by chromstaR and considered only expressed TFs. We used the R package TFBSTools ^98^ to compute the similarities between any two binding motifs based on their Pearson correlation. We clustered together TFs with a similarity higher than 0.9, obtaining 255 distinct unique TF clusters. To investigate TF co-binding, we joined chromatin regions containing TF binding instances less than 500 bp apart, resulting in a set of chromatin regions with variable sizes (up to 3 kbp) concentrating TF activity during the entire time course, as described in ^27^. We performed a hypergeometric test for each pairwise co-binding possibility for each time point and adjusted p-values for multiple testing using Bonferroni correction.

### TRANSCRIPTION FACTOR NETWORKS

We built TF hierarchical networks as previously described in Field 27, 95, focusing on transcriptionally upregulated TFs from the OIS gene module to evaluate the TF hierarchical and temporal activity. Briefly, each identified TF is represented as a node, connected by directed edges representing co-binding events along the chromatin and during the entire time course. An edge with TF A as the source and TF B as the target indicates that TF A binds to the same chromatin regions as TF B at the same or at a previous time point. For each TF, we computed their normalized total number of bound regions and dynamicity and represented those properties in the networks as node size and node color, respectively. Each edge is associated with a weight in the interval [0, 1], representing the fraction of TF B binding instances previously occupied by TF A. We filtered the networks for edges with a weight higher than 0.15 and performed a transitive reduction to simplify the obtained networks while keeping essential topological features. Nodes included in the same strongly connected component (SCC), i.e., connected both by incoming and outgoing paths, were merged into a single node. We represented the identified transcriptomic module distribution for TFs included in the same SCC as Doughnut Charts. We used the R package igraph to represent the TF networks and exported them from the R environment to the Cytoscape software using the R package Rcy3 ^99^ and the CyREST API ^100^.

### NORMAL COLON AND COLORECTAL CANCER GENE EXPRESSION AND CHROMATIN ACCESSIBILITY INTEGRATION

Fastq files of normal colon DNA-seq datasets from the ENCODE Consortium ^101, 102^ with identifiers ENCSR088CMZ, ENCSR340MRJ, ENCSR276ITP, ENCSR923JYH, and ENCSR760QZM (all generated by the John Stamatoyannopoulos laboratory) were downloaded from SRA and aligned to the GRCh38.d1.vd1 version of the human genome using bowtie2 with the local alignment option. Alignments were preprocessed as described above for ChIP-seq and ATAC-seq datasets. Alignments of 38 ATAC-seq datasets of colorectal cancer tumor samples available at the Cancer Genome Atlas (TCGA) were downloaded. Differential accessibility and clustering were performed using DESeq2 and WGCNA. Footprinting and differential binding activity were performed using HINT-ATAC as described above. Senescence scars in CRC were identified with bedtools *intersect* using the GRCh38.d1.vd1 coordinates of senescence scars identified in GM21 and the CRC-specific differentially accessible modules as input. Where indicated in the main text, integration with differential expression data from CRC was performed by annotating POU2F2 footprints and senescence scars to the nearest gene, and genes with an absolute log2 fold change of 0.29 were considered. Differential expression data was obtained from the Xena Differential Gene Expression Analysis Pipeline comparing 631 primary tumors to 101 solid normal tissue biopsies ^103^. Functional characterization of POU2F2 OIS escape and senescence scar CRC-specific gene signatures was performed with clusterProfiler using MSigDB and KEGG pathway gene sets. The results shown here are in whole or part, based upon data generated by the TCGA Research Network: https://www.cancer.gov/tcga.

### GSEA

GSEA was performed on transcriptomes (GSE32323 and GSE35896) of patients with CRC phenotypically classified as in ^69, 70^ using GSEA GUI version 4.2.2. Probe sets were collapsed to the gene level using the correlation-based approach^104^, whereby the correlation of probe sets representing the same gene was computed to decide whether to average probe sets (c > 0.2) or to use the probe set with the highest average expression across samples (c ≤ 0.2). Probe sets without known annotation were removed. The signal-to-noise ratio (μA – μB)/(σA + σB) (μ represents the mean, σ the standard deviation) was used as a ranking metric and statistics based on gene set permutations. For POU2F2 RNA interference experiments, the list of predicted POU2F2 target genes defined by ATAC-seq footprinting was used as input to determine positively and negatively regulated POU2F2 target genes. The labels ‘pLKO’ and ‘shPOU2F2’ were used to define the phenotypes according to the shRNA treatment, and genes were ranked from high to low expression relative to ‘pLKO’. GSEA was performed using the log2-ratio-of-classes as a ranking metric and statistics calculated based on gene set permutations. Positively and negatively regulated genes were selected by subsetting target genes at ranking metric of >= 300 and <= -300, respectively.

### EDU INCORPORATION AND POPULATION DOUBLINGS

Growth curves were generated by counting the cells with a hematocytometer and using the formula PD=3.32*[log(N final)-log(N initial)], where N final is the number of cells recovered from the dish and N initial is the number of cells initially seeded at each plate. To measure EdU incorporation, cells were labeled with 10μM EdU for 48 hrs, and EdU positive cells were detected using the ClickiT Plus EdU Alexa Fluor 488 or 594 imaging kit (Thermo Fisher Scientific, Waltham, MA) according to the manufacturer’s instructions. Images were acquired using a Zeiss Axio Observer Z1, an AxioCam MRm camera (Zeiss, Jena, Germany), and Zen Blue 2.3 software (Zeiss, Jena, Germany). For EdU incorporation, 3 to 5 images per coverslip were imaged at 10x magnification, and positive signals and a total number of cells were manually counted with Zen Blue 2.3 software.

### WESTERN BLOTTING

Protein extracts were prepared in 1x CHAPS Lysis buffer (Millipore-Sigma, Saint Louis, MO) containing 1:100 Halt protease and phosphatase inhibitor cocktail (Thermo Fisher Scientific, Waltham, MA). A total of 10 to 20 μg of whole-cell lysates were resolved on 4-15% SDS/PAGE precast gels (Bio-rad, Hercules, CA), and proteins were transferred to nitrocellulose membranes using the Trans-Blot Turbo Transfer System (Bio-rad, Hercules, CA) and probed as described ^10^. The following primary antibodies and dilutions were used: anti-Ras (BD Transduction Laboratories, San Jose, CA; Cat no. 610001; 1:1000 dilution), POU2F2 (Santa Cruz Biotechnology, Dallas TX; Cat no. sc-377475-X; 1:500 dilution) and γ-Tubulin (Millipore-Sigma, Saint Louis, MO; Cat no. T6557; 1:1000 dilution). HRP-conjugated goat anti-rabbit (PerkinElmer, Waltham, MA; Cat no. NEF812001EA; 1:6000 dilution) or anti-mouse (Cell Signalling, Danvers, MA; Cat no. 70765; 1:3000 dilution) antibodies were used as secondary antibodies. Densitometric analysis of CDKN2A/p16^Ink4a^ in shPOU2F2 cells is presented as fold increase compared to control plkO.1 EV cells. γ-tubulin is used as loading control. Bands were visualized using Clarity Max ECL system (Bio-Rad, USA) and were quantified using Image Studio Lite Ver. 5.2.

### REAL-TIME QUANTITIVE POLYMERASE CHAIN REACTION

Total RNA was isolated from cells with RNeasy Mini kit (Qiagen, Germantown, MD) or Nucleospin RNA Plus XS (Macherey-Nagel, Allentown, PA) according to the manufacturer’s protocol. As per the manufacturer’s instructions, reverse transcription of 500 ng of total RNA was carried out using the iScript cDNA synthesis kit (Bio-rad, Hercules, CA). Quantitative real-time PCR (qPCR) was performed using specific primers (listed below) with the iTaq Universal SYBR Green Supermix (Bio-rad, Hercules, CA) on a CFX96 Touch Real-Time PCR detection system (Bio-rad, Hercules, CA). Samples were analyzed in technical triplicates, and GADPH levels were used for normalization. The following Qiagen QuantiTect Primers were used: GADPH (QT00079247), HRAS (QT00008799), IL1B (QT00021385), IL8 (QT00000322), CDKN2A (QT00089964), CDKN1A (QT00062090), CCNA2 (QT00014798) and POU2F2 (QT00087290) (Qiagen, Germantown, MD).

### TISSUE PROCESSING

Tissue samples were collected from patients undergoing partial or complete colectomy surgery with the approval of the local IRB committee. Following histopathological analysis by the surgical pathologist, tissue considered excess was provided. Samples were transported on ice and immediately cryopreserved in OCT compound (Sakura, Torrence, CA) using 2- methylbutane cooled with liquid nitrogen. Samples were stored at –80 °C until ready for use.

The tissue was cut into 8 µm sections using a cryomicrotome (Leica, Dublin, OH) set to -16 °C mounting block temperature and -20 °C air temperature in the chamber and affixed to room temperature positively charged microscope slides (Fisher, Waltham, MA). Slides were then placed on dry ice for transport to storage at –80 °C until staining. H&E stain was performed by serial submission in hematoxylin, mild-acid differentiator, bluing solution, and eosin with wash steps after each, followed by dehydration with ethanol and xylenes before mounting coverslips with Cytoseal (Thermo Fisher Scientific, Waltham, MA). All tissue samples were collected from NJMS-University Hospital with the approval of the local IRB committee.

### IMMUNOHISTOCHEMISTRY

Tissue samples were brought to room temperature and outlined in a hydrophobic pen. PBS was used to dissolve excess OCT, and samples were then fixed in 4% PFA for 3 minutes.

Immunohistochemistry was performed in duplicate for each sample per manufacturer’s instructions using Mouse and Rabbit specific HRP/DAB (ABC) Detection IHC Kits (Abcam, Cambridge, UK) with TBST used for wash buffer. Briefly, slides were blocked with peroxide block, then protein block for ten minutes each, incubated with primary antibody for one hour at room temperature,e followed by incubation with biotinylated goat anti-mouse or anti-rabbit secondary and streptavidin HRP for ten minutes each. DAB chromogen/substrate mixture was applied for 1 minute, and samples were given a final rinse in deionized water. The primary antibodies used were monoclonal mouse anti-POU2F2 (Santa Cruz Biotechnology, Dallas, TX; Cat no. sc-377475-X). One duplicate of each stain was counterstained with hematoxylin and ammonium bluing solution, then all samples were dehydrated with ethanol and xylenes and coverslips mounted with Cytoseal (Thermo Fisher Scientific, Waltham, MA). Tissues were visualized on a Zeiss AX10 microscope, images acquired in AxioVision version 4.7.1, and processed on ImageJ version 1.51.

### TELOMERASE ACTIVITY

Telomerase activity was measured by the TRAPeze RT Telomerase detection Kit (Millipore- Sigma, Saint Louis, MO), using PCR and Amplifluor primers according to the manufacturer’s instructions. Pelleted cells were resuspended in CHAPS lysis buffer, incubated on ice for 30 min, and spun to collect the supernatant. A total of 400 ng of protein lysate was used to detect telomerase activity using the Real-Time PCR system (Bio-Rad). Quantitative values of the telomerase activity were obtained from a standard curve generated by dilutions of the control template provided by the kit.

### QUANTIFICATION OF TELOMERE DYSFUNCTION-INDUCED DNA DAMAGE FOCI

Images were acquired using a Zeiss Axiovert 200M microscope, an AxioCamMRm camera (Zeiss, Jena, Germany), and AxioVision software (Version 4.6.3; Zeiss). To analyze and quantitate colocalization between telomere signals and DSB foci, images were acquired as z-series (7 to 11 images, 0.3-μm optical slices) using a motorized stage, a 63×/1.4 oil immersion lens, and an ApoTome (Zeiss, Jena, Germany). ApoTome microscopy eliminates out-of-focus light and generates shallow focal planes (0.5 μM using a 63× oil objective). Random images of at least 100 cells were acquired using a 63×/1.4 oil immersion lens to quantitate DDR foci.

### COLONY FORMATION ASSAY

5,000 cells that entered into and escaped from OIS and time-matched empty vector controls were mixed with a final concentration of 0.3% agar and DMEM medium containing 10% FBS and plated in triplicates onto six-well dishes containing a basal layer of 0.8% agar containing DMEM medium and 10% FBS. The plates were kept in a humidified incubator at 37 °C with 5% CO2 and 2% Oxygen for five weeks. Fresh medium was added every four days. After five weeks, digital images of 20 random fields at 10x magnification were acquired using Zeiss Axio Observer Z1 with an attached AxioCam MRm camera (Zeiss). The number and size of colonies were counted manually using Zen Blue 2.3 software (Zeiss).

---

**Figure S1.**
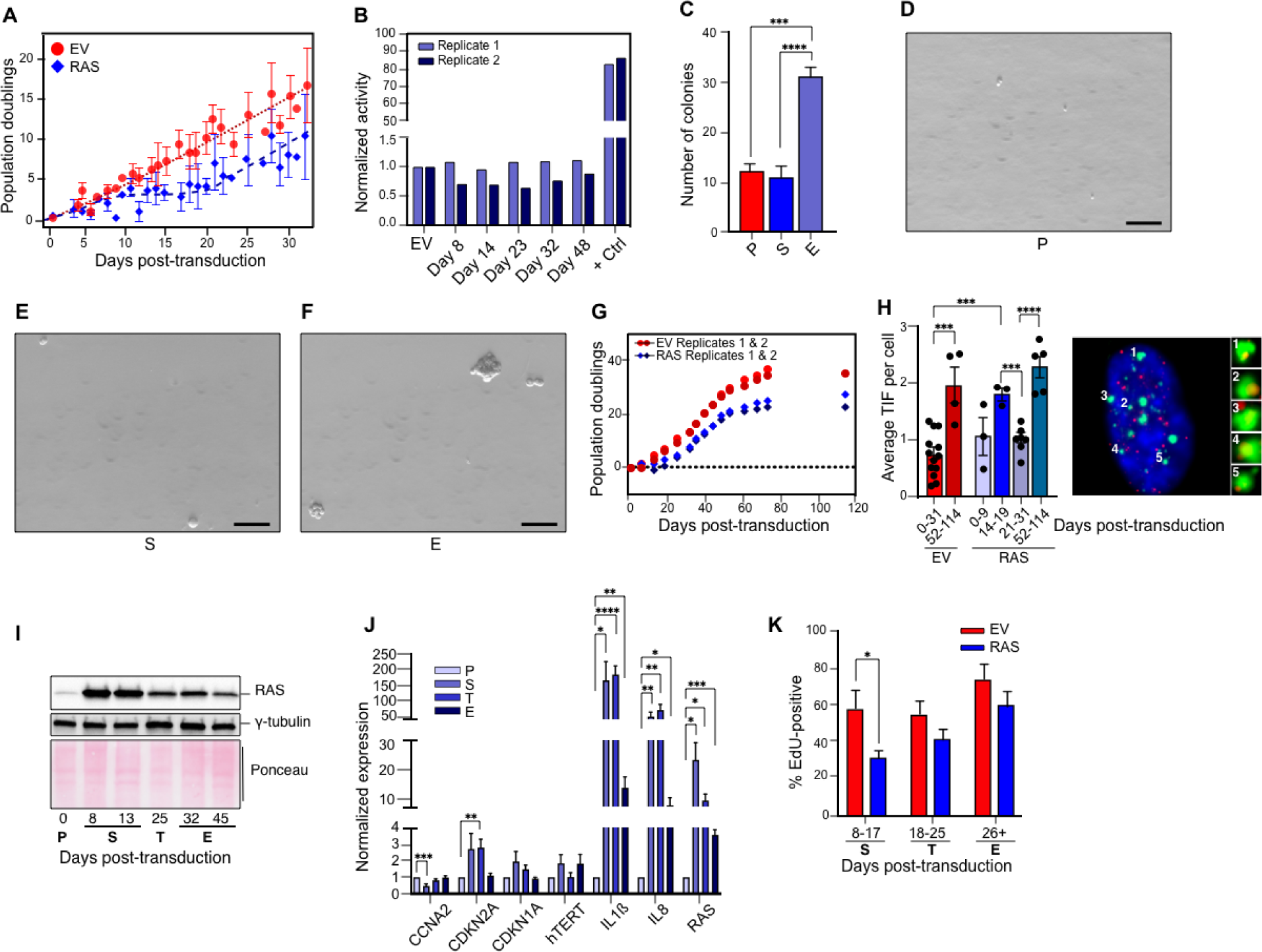
Molecular characterization of OIS escape. **(A)** Population doubling curve of GM21 fibroblasts transduced with an empty vector (red circles) or a vector expressing H-RAS^G12V^ (blue rhombuses). Fourteen (EV, empty vector) and fifteen (RAS) biologically independent time-series experiments are represented in the plot. For time points with repeated measures, the bars indicate the standard error of the mean. Dotted and hashed lines are illustrative and do not represent any statistical procedure on the data. **(B)** Quantification of telomerase activity in GM21 fibroblasts transduced with an empty vector and at the indicated time points after overexpression of H-RAS^G12V^. A positive control is included. Two biologically independent experiments are shown. **(C)** Quantification of colony formation in soft agar in GM21 fibroblasts that escaped OIS and time-matched empty vector controls (day 32 post-transduction). Abortive colonies were defined as >2 cells. Bars and whiskers show the average and standard error of the mean of n=3 (EV) and n=6 (S and P). Statistical significance was calculated with a two- tailed, unpaired *t*-test for the pairwise comparisons. ***, *p* = 0.0006; ****, *p* < 0.0001. **(D-F)** Representative micrographs of colonies of GM21 fibroblasts at P, S, and E. Scale bar, 50 µm. **(G)** Population doubling curve of GM21 fibroblasts transduced with an empty vector (red circles) or a vector expressing H-RAS^G12V^ (blue rhombuses). Two biologically independent experiments are shown. **(H)** Quantification of telomere dysfunction-induced DNA damage foci (TIF) in control empty vector and H-RAS^G12V^-overexpressing GM21 fibroblasts at the indicated time points. Circles represent the average TIF per cell per time point within the indicated ranges. Graphs show the average +/- standard error of the mean (s.e.m) in three biologically independent experiments. A representative micrograph showing the presence of TIF is shown. Statistical significance was calculated with a one-tailed Student’s *t*-test for each pairwise comparison indicated by the brackets. ***, *p* = 0.0008; ****, *p* < 0.0001. **(I)** Representative Western analysis of oncogenic RAS expression in an OIS escape time course. **(J)** RT-qPCR profiling of selected senescence biomarkers throughout the stages of escape from OIS. Statistical significance was calculated with a one-tailed Student’s *t*-test, with P as reference for each pairwise comparison, as shown. *, *p* < 0.05; **, *p* < 0.005; ***, *p* < 0.001; ****, *p* < 0.0001. Graphs show the average +/- standard error of the mean (s.e.m) of at least three biologically independent experiments. **(K)** Quantification of EdU incorporation at the indicated stages during entry into and escape from OIS. Bars represent the average of at least three biologically independent experiments, and whiskers show the standard error of the mean. Statistical significance was calculated with an unpaired *t*-test with a single pooled variance and Holm-Sidak multiple test correction. *, *p* = 0.048.

**Figure S2.**
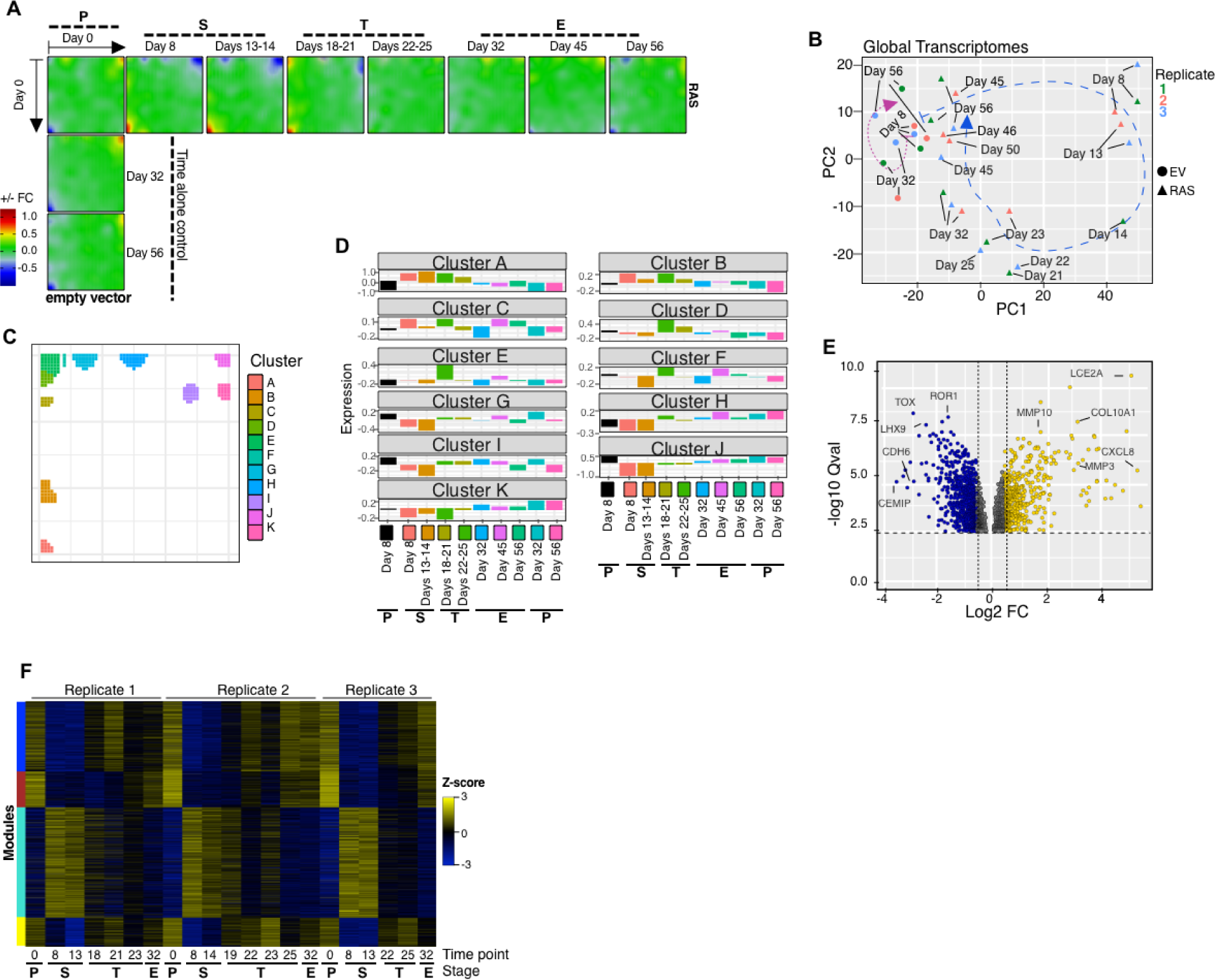
Transcriptome and epigenome dynamics of OIS escape. **(A)** Averaged SOMs of transcriptomes of three biologically independent time-series experiments of GM21 fibroblasts undergoing escape from OIS (horizontal) and a time-matched empty vector control (vertical) expressed as logarithmic fold-change measured until day 56 after H-RAS^G12V^ transduction. The transcriptional landscape of each stage of OIS escape is highlighted. **(B)** Principal component analysis projection plots showing the transcriptional trajectories of global transcriptomes of three biologically independent OIS escape time-series measured until day 56 after H-RAS^G12V^ transduction. **(C)** D-cluster projection showing regions of SOMs with significant changes in expression throughout escape from OIS (see regions of SOMs in (**A**) that change in color through time). **(D)** Expression of D-cluster metagenes in GM21 fibroblasts entering into and escaping from OIS and empty vector controls. The histograms show the change in expression of metagenes in clusters identified in (**C**) (top of panels) and are color-coded according to condition and time point (bottom of panels). **(E)** Volcano plot collapsing all differentially expressed genes in GM21 fibroblasts entering into and escaping from OIS in at least one-time point relative to empty vector control cells. **(F)** Heatmap showing the dynamic expression profiles of gene expression modules in individual time-series experiments.

**Figure S3.**
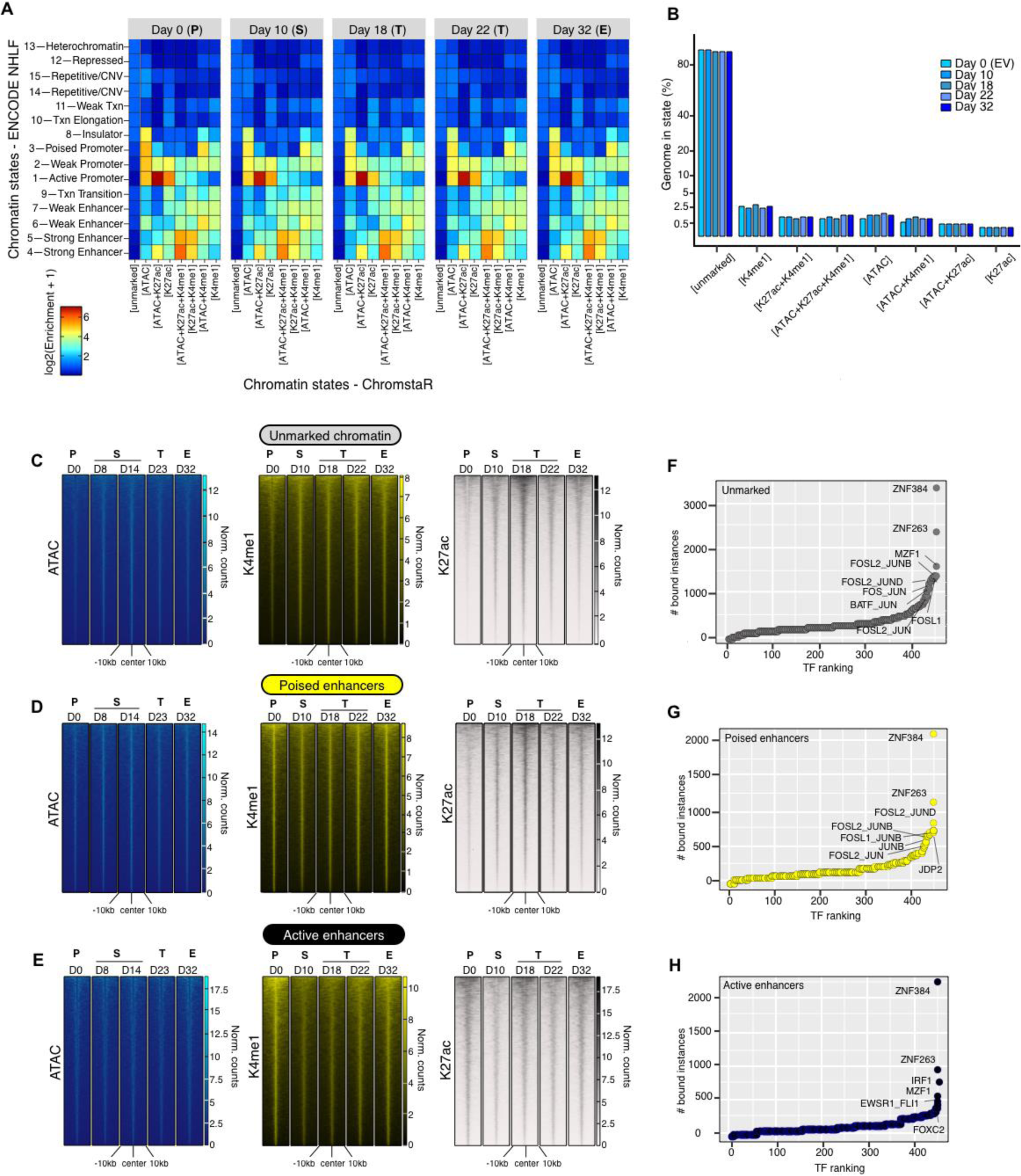
Transcriptome and epigenome dynamics of OIS escape. **(A)** Enrichment heatmap showing the combinations of H3K4me1, H3K27ac ChIP-seq, and ATAC-seq signals with reference chromatin states in normal human lung fibroblasts from the ENCODE project. Txn, transcription; CNV, copy number variation. **(B)** Histograms showing the percentage of the genome enriched in the indicated combinations of H3K4me1, H3K27ac ChIP-seq, and ATAC-seq signals at the indicated time points in GM21 fibroblasts overexpressing oncogenic RAS. **(C- E)** Genome-wide signal intensity evolution heatmaps of ATAC-seq, H3K4me1, and H3K27ac in unmarked chromatin (**C**), poised (**D**), and active (**E**) enhancers at day 0 in GM21 fibroblasts undergoing escape from OIS. **(F-H)** Rank plots showing the summed binding instances of TFs in unmarked chromatin (**F**), poised (**G**), and active (**H**) enhancers at day 0. Data in **A**-**E** was computed from two independent immunoprecipitations per time-point per histone modification from pooled chromatin from 10 biologically independent experiments. Data in **F**-**H** is the average of two biologically independent ATAC-seq time series.

**Figure S4.**
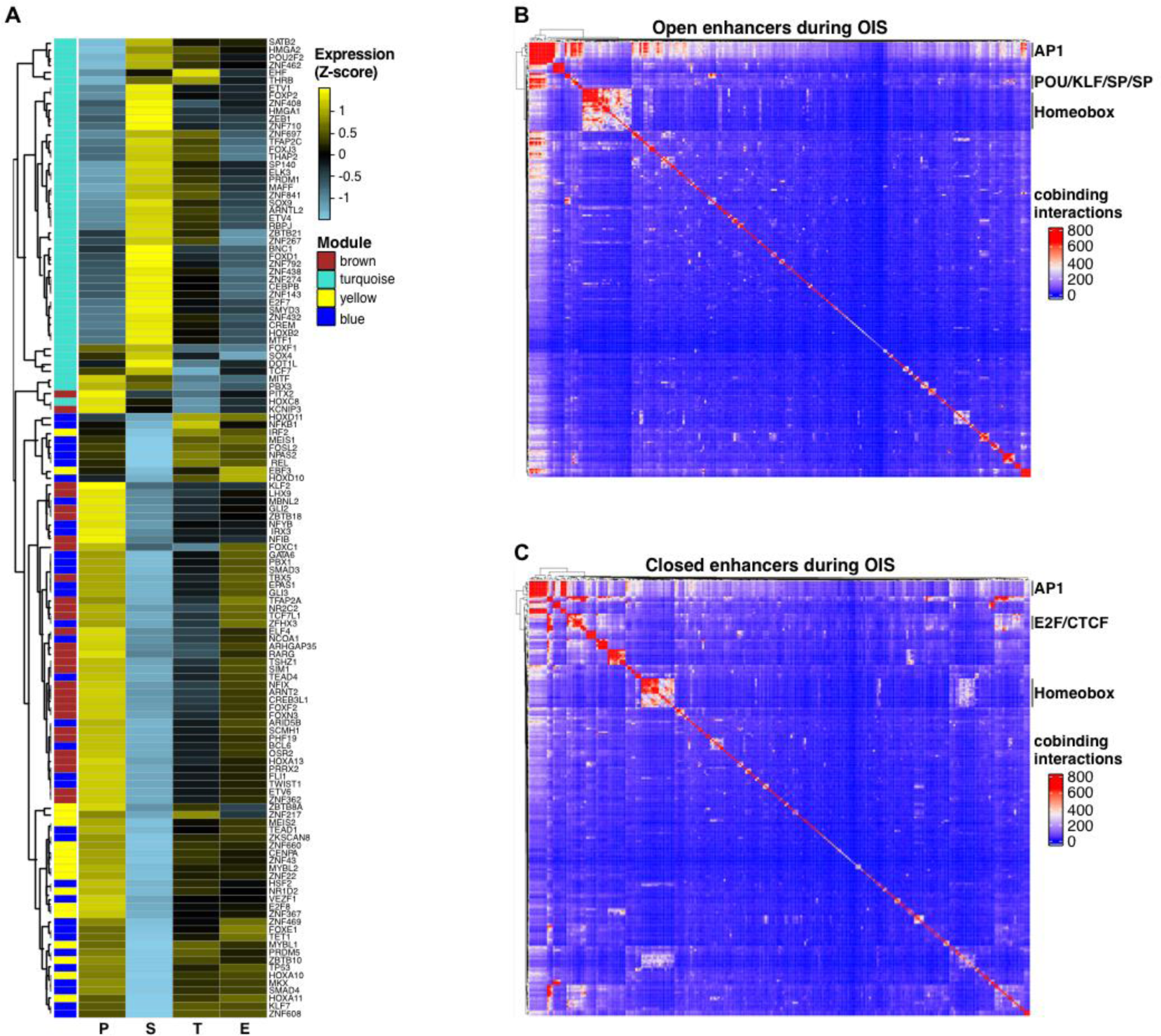
Organized waves of TF activity define OIS escape. **(A)** Heatmap showing the dynamic expression profile of TFs within each transcriptomic module throughout each stage as cells enter and escape from OIS. **(B,C)** TF co-binding matrices at open (**B**) and closed (**C**) enhancers during OIS in GM21 fibroblasts entering and exiting from OIS. All binding instances across time points were collapsed onto the matrix and clustered by Ward’s aggregation criterion. The matrices are color-coded based on the number of co-binding interactions computed per TF (row sum). TF expression (**A**) and footprinting (**B**,**C**) were performed on pooled transcriptome (n=3) and ATAC-seq (n=2) data sets from biologically independent time- series experiments.

**Figure S5.**
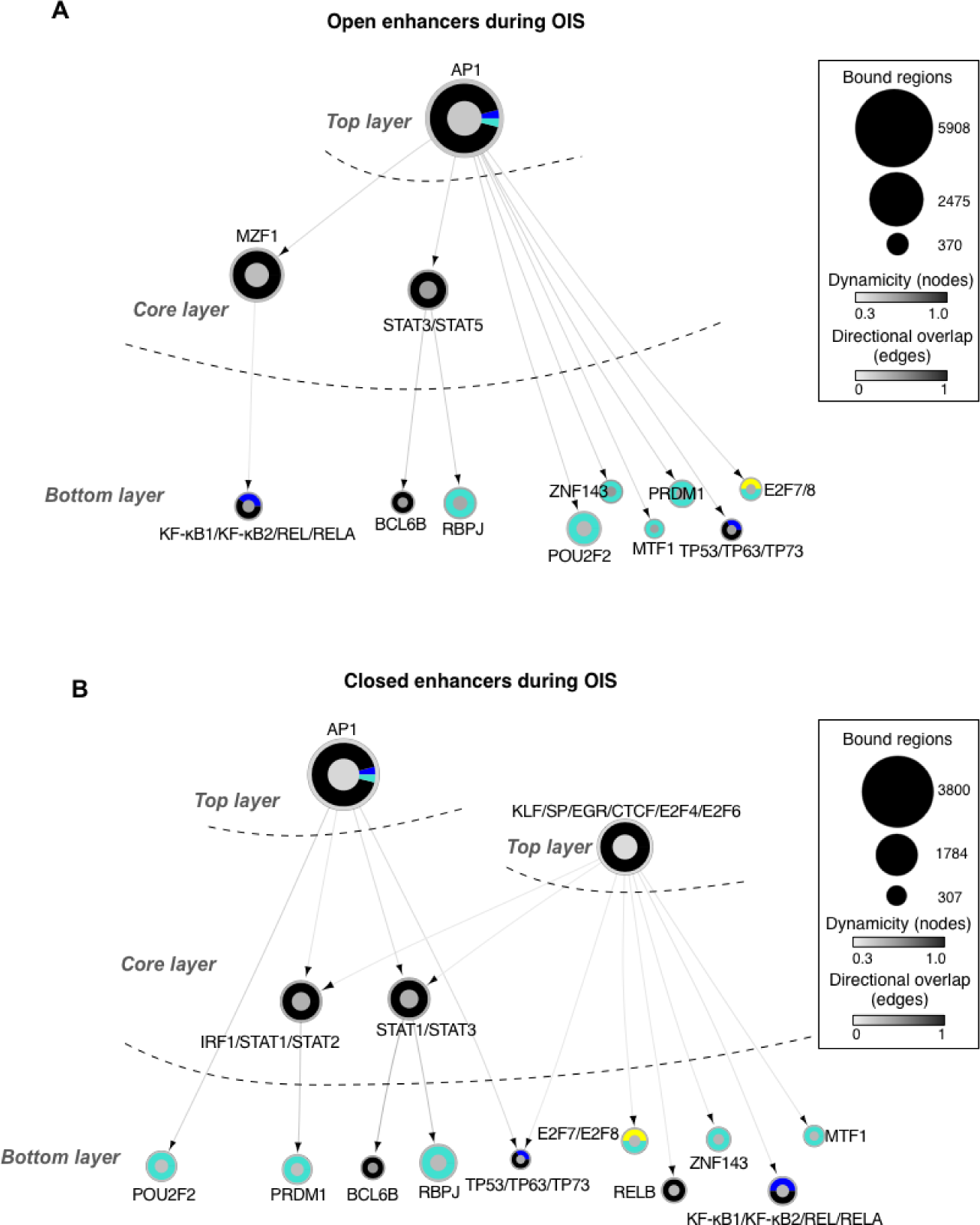
Hierarchical TF networks define transcriptional dynamics of OIS escape. (A,B) Effector TF networks during OIS at open (**a**) and closed (**b**) enhancers. TFs (nodes) are represented as circles. Oriented edges (arrows) connecting nodes indicate that at least 15% of the regions bound by a given TF in the bottom and core layers were bound by the interacting TF in the core and top layers, respectively, at the same or previous time points. Strongly connected components (SCCs) are represented as a single node to facilitate visualization. The fill color of the node’s inner circle is based on the normalized dynamicity of TFs. The fill color of the outer ring indicates whether the TF is constitutively expressed (black) or belongs to a transcriptomic module (blue, brown, turquoise, and yellow from Figure 1e). The node’s size is proportional to the bound regions by a given TF. Each network has three layers: i) top layer with no incoming edges, ii) core layer with incoming and outgoing edges, and iii) bottom layer with no outgoing edges. Networks were generated from pooled ATAC-seq data sets from two biologically independent time series.

**Figure S6.**
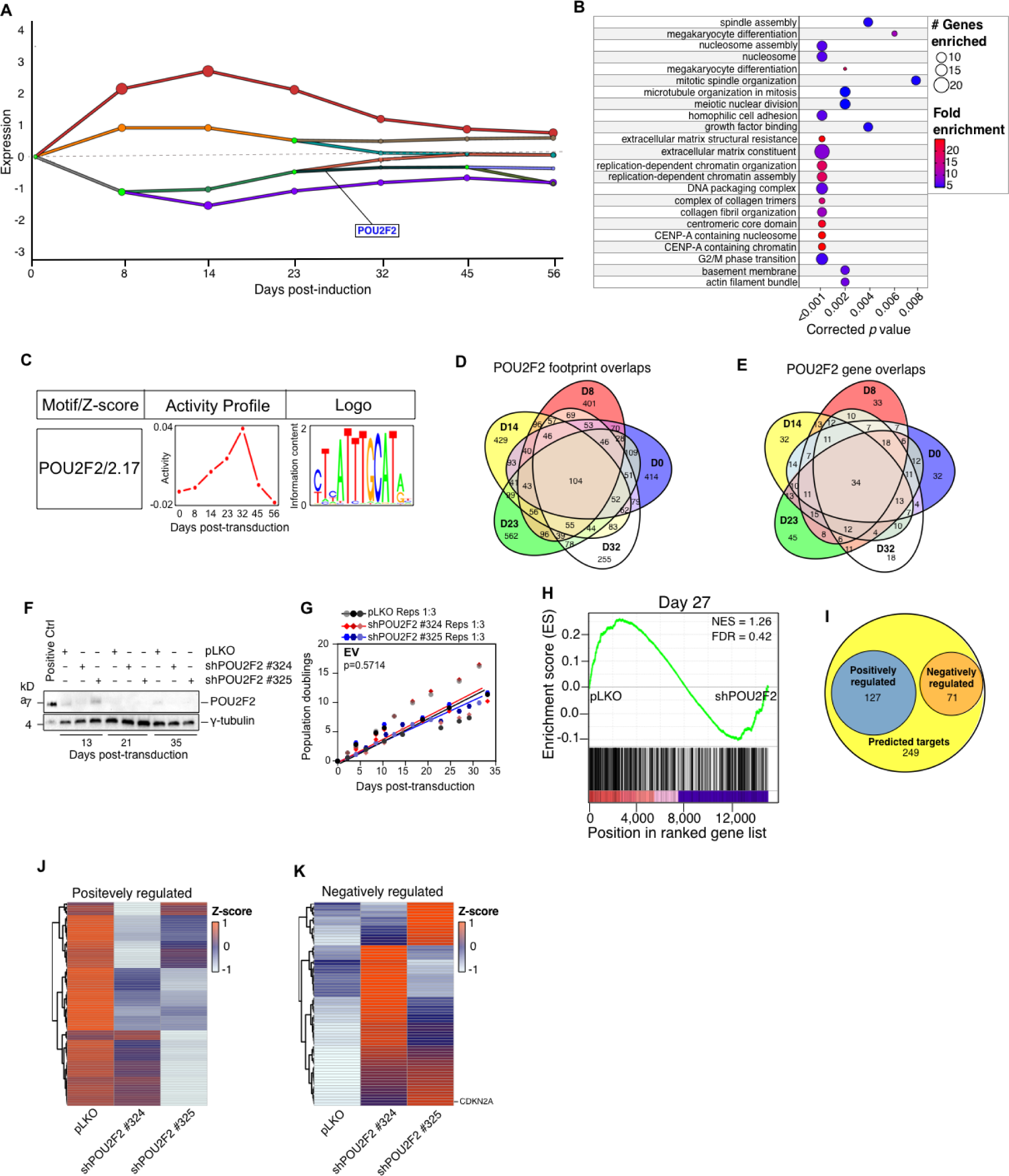
POU2F2 promotes OIS escape. **(A)** Dynamic regulatory event miner (DREM) gene expression trajectory map showing paths of co-expressed genes. POU2F2 is transcriptionally upregulated and predicted to regulate the expression of genes in the indicated path. **(B)** Gene Ontology (GO) overrepresentation analysis of genes in the POU2F2-regulated expression path. Circle fill is color-coded according to the fold enrichment of the genes in the path within each GO gene set (rows). Circle size is proportional to the number of genes in each GO gene set. The corrected *p* values (bottom) were calculated by randomization tests using 500 samples of the analyzed gene set. **(C)** Summarized output from the Integrated System for Motif Activity Response Analysis (ISMARA) showing increased POU2F2 binding activity at promoters of genes as cells enter into and escape from OIS. Results in (**A-C**) were obtained using transcriptome data from three biologically independent experiments. **(D,E)** Intersections and disjunctive unions of POU2F2 binding (**D**) and gene targets (**E**) at each time point during the escape from OIS. **(F)** Representative Western blot analysis of POU2F2 expression in GM21 fibroblasts constitutively expressing non-targeting and POU2F2 targeting shRNAs and transduced with empty vector control. The positive control is from GM21 fibroblasts overexpressing H-RAS^G12V^ on day 19. One of four biologically independent experiments is shown. **(G)** Population doubling curve of GM21 fibroblasts constitutively expressing non- targeting and POU2F2 targeting shRNAs and transduced with empty vector control. The projected lines and *p-value* were calculated using three biologically independent experiments and simple linear regression. **(H)** GSEA showing normalized enrichment score (NES) plots and FDR values for POU2F2 predicted target genes (**5C**) in transcriptomes of pLKO and shPOU2F2-expressing cells at day 27 after RAS overexpression. Statistical evaluation of GSEA results was based on a nonparametric Kolmogorov–Smirnov test. **(I)** Intersections of positively and negatively regulated POU2F2 target genes within the predicted gene set defined in (**5C**). **(J,K)** Expression heatmap of positively (k) and negatively (l) POU2F2-regulated genes.

**Figure S7.**
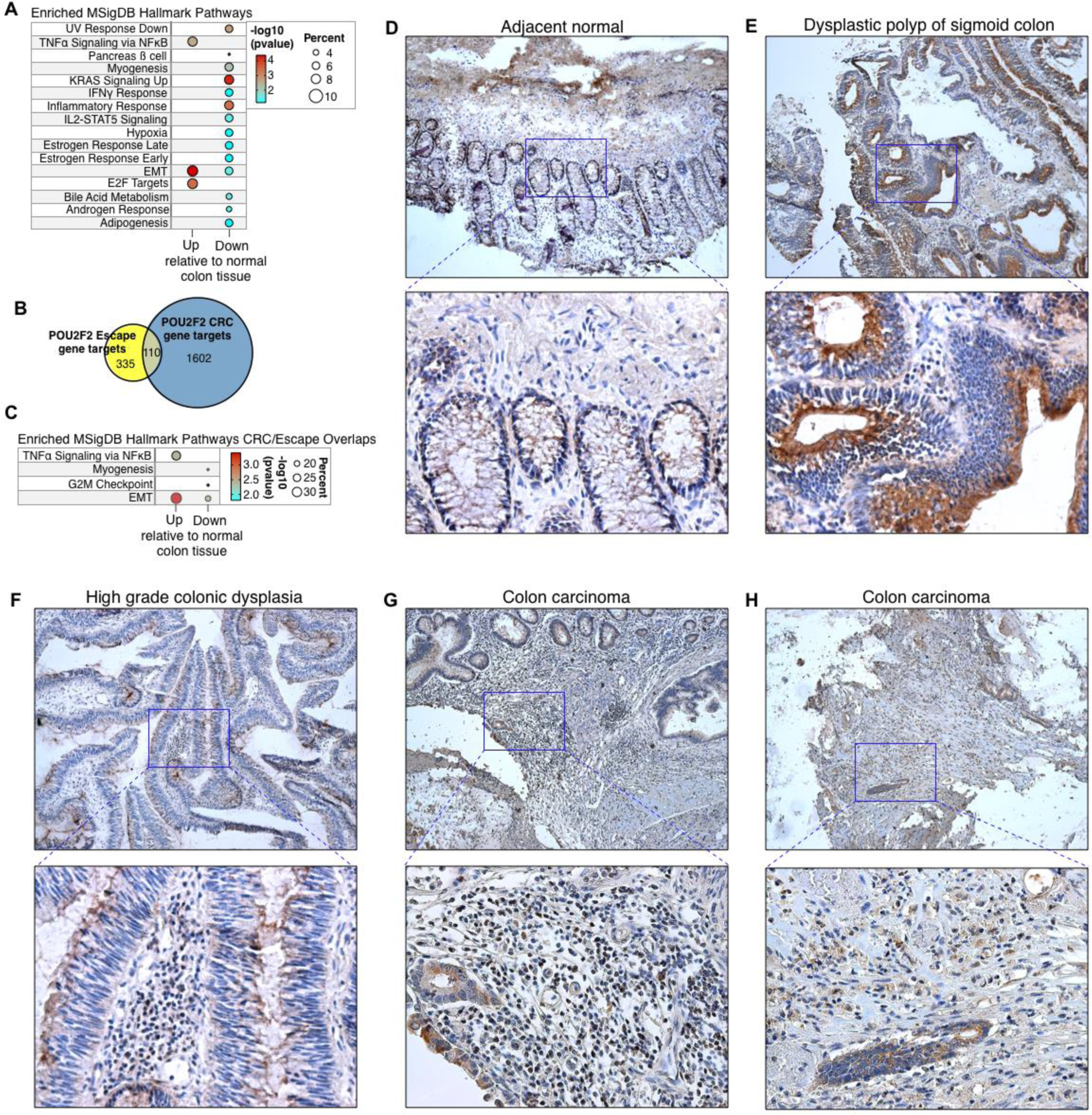
POU2F2 drives an inflammatory gene expression program in colorectal cancer. **(A)** Functional over-representation analysis map showing significant associations of the MSigDB Hallmark gene sets with the POU2F2 CRC gene targets in the differentially expressed genes in CRC relative to normal colon tissue. Circle fill is color-coded according to the FDR- corrected *p-value* from a hypergeometric distribution test. Circle size is proportional to the percentage of genes in each MSigDB gene set found within each gene group. POU2F2 gene targets in CRC were identified using gene expression data from 101 normal colon and 631 primary tumors deposited in the TCGA using the Xena Differential Gene Expression Analysis Pipeline. **(B)** Intersections and disjunctive unions of POU2F2 gene targets in GM21 fibroblasts escaped from OIS and CRC. **(C)** Functional over-representation analysis map showing significant associations of the MSigDB Hallmark gene sets with the POU2F2 gene targets common to cells that had escaped from OIS and CRCs, both overexpressed and downregulated relative to normal colon tissue. Circle fill is color-coded according to the FDR-corrected *p-value* from a hypergeometric distribution test. Circle size is proportional to the percentage of genes in each MSigDB gene set found within each gene group. **D-H** Representative micrographs at 10X (top) and 40X (bottom) magnification of POU2F2 expression levels in adjacent normal tissue (**D**), a dysplastic polyp of the sigmoid colon (**E**), a high-grade colonic dysplasia (**F**), and colon carcinoma (**G,H**) from three patients that underwent colectomy procedures. Sections were stained for Hematoxylin to identify nuclei.

**Figure S8.**
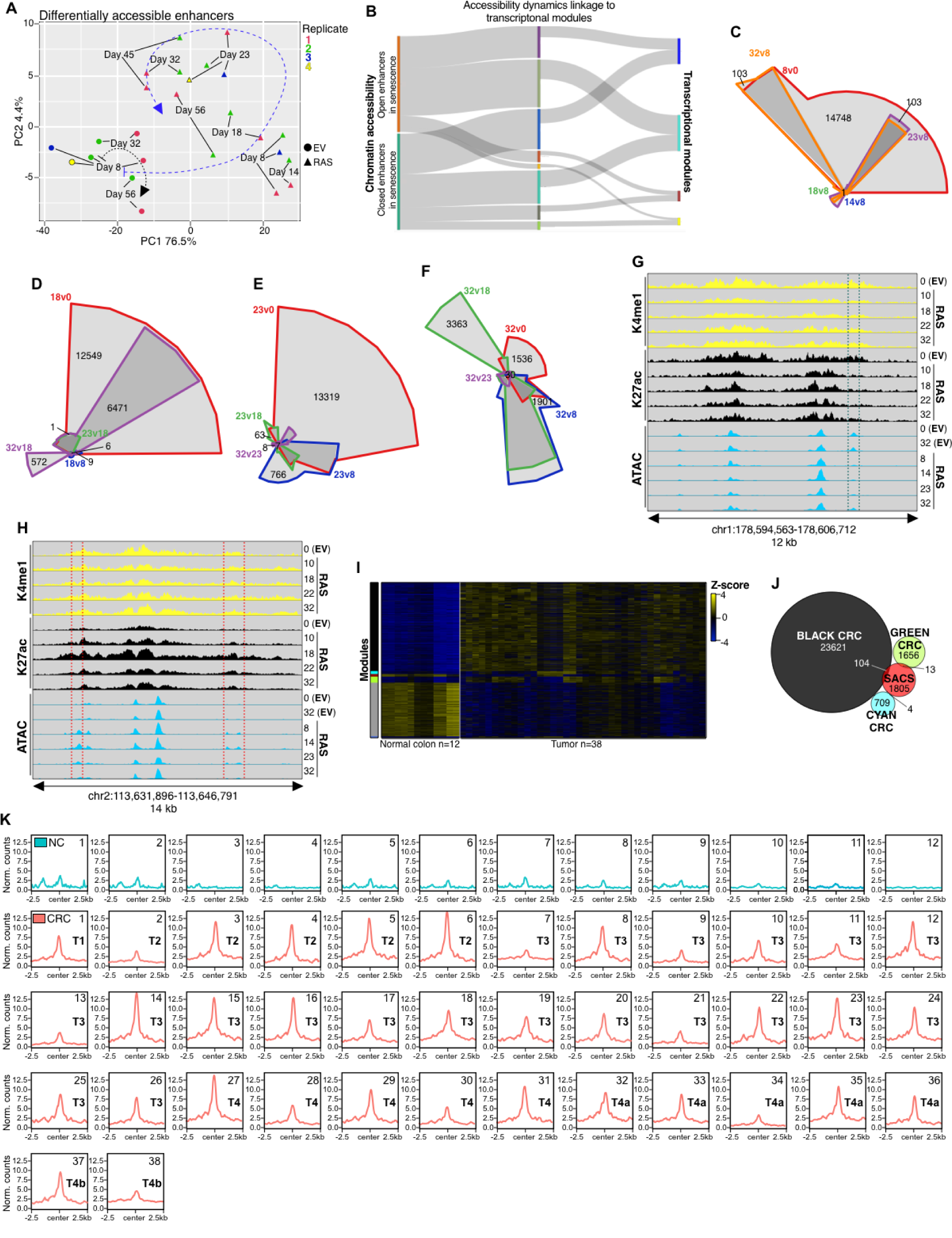
SACS define post-senescent cells and are present in colorectal cancer. **(A)** Principal component analysis projection plots showing the trajectories of differentially accessible chromatin regions of the indicated biologically independent replicates per time point of GM21 fibroblasts overexpressing RAS^G12V^ and empty vector controls. **(B)** River plots showing the relationship between differentially accessible chromatin modules and gene expression modules. Data were computed from pooled ATAC-seq and transcriptome data sets from two and three biological independent time-series experiments. **(C-F)** Chow-Ruskey diagram showing intersections and disjunctive unions of differentially accessible chromatin regions at each time point measured relative to days 8 (**C**), 18 (**D**), 23 (**E**), and 32 (**F**). **(G,H)** Representative genome browser screenshots highlighting the normalized signal evolution of H3K4me1, H3K27ac, and ATAC-seq at closing (**G**) and opening (**H**) senescence scars. Data in **C**-**F** was computed from the average of two biologically independent ATAC-seq time series. Data in **G**,**H** shows the average signal of two independent immunoprecipitations per time-point per histone modification from pooled chromatin from 10 biologically independent experiments and two biologically independent ATAC-seq time series. **(I)** Heatmap showing differentially accessible chromatin modules in normal human colon and CRC. Signal intensity is represented as row Z-scores. *N* for normal colon and tumors is shown. **(J)** Intersections and disjunctive unions of CRC-specific accessible chromatin and open senescence scars. **(K)** Individually normalized meta profiles of senescence scars in normal colon and CRC. The AJCC TNM stage is indicated in tumor samples.

